# Phosphate and Carbonate in the Biomineralization of Chicken Eggshells and the Increase in Eggshell Thickness through Nanodroplet Addition

**DOI:** 10.64898/2026.02.23.707050

**Authors:** Antonio Valadão Cardoso, Rodrigo Novaes Ferreira, Maria Sylvia Dantas, Leonardo Humberto Rezende dos Santos, Ana Paula Gomes, Lara Luz de Assis

## Abstract

The presence of hydroxyapatite (HAp) in the Cuticle of laying hen eggshells was investigated through an extensive and detailed study combining scanning electron microscopy (SEM) coupled with energy-dispersive spectroscopy (EDS), as well as micro-Raman, micro-FTIR, X-ray diffraction (XRD), and thermogravimetric analysis (TG). Additionally, Raman and FTIR spectra of Cuticle HAp were compared with those obtained from the internal surface of chicken femur fragments and from a bovine HAp sample.

Examination of the same region by SEM in SE (topographic) and BSE (subsurface compositional contrast) modes revealed unidirectional (nanofibrous) calcite growth within the Vertical Layer (VL) and Palisade Layer (PL), which together constitute nearly the entire eggshell thickness. Shell thickening in the VL and PL layers appears to proceed via an additive mechanism characterized by the successive deposition of nanodroplets containing, according to our hypothesis, the mineral phase, water, and organic components. This multiphasic system generates lamellae that progressively increase in thickness through the continuous incorporation of new nanodroplets onto the pre-existing surface. This additive nanodroplet-mediated growth contributes to understanding how micropores form in the PL and VL.

Biomineralization via an additive mechanism is strongly supported by the presence of nano-hemispheres attached to growing lamellae in the VL and PL. Statistical analyses corroborate the relationship between the diameter of Cuticle nanospheres and that of nano-hemispheres in the Vertical Layer. Fractures observed in the VL indicate structural continuity between the Cuticle and the Vertical Layer, suggesting that additive growth involves a continuous supply of HAp — possibly across the entire uterine surface — which, through a yet undescribed mechanism, dissolves and/or transforms calcium phosphate nanospheres into calcium carbonate nanofibers.

## 1. INTRODUCTION

Laying hens accomplish the remarkable feat of biomineralizing an eggshell within a period of 18–20 hours, producing a shell approximately 0.5 mm thick (1). This remarkable capacity occurs during the laying period of the hen and may last for 2 to 3 years depending on the living conditions of the bird (2). On average, a hen lays 500 eggs during its active life, which corresponds to an accumulated thickness of nearly half a meter of calcite (CaCO₃) biomineralization.

Over the past 30 years, research on the biomineralization of the eggshell of laying hens (*Gallus gallus domesticus*) has advanced considerably. The combination of Molecular Biology techniques with theories and methods from Materials Science has proven highly fruitful in advancing our understanding of the eggshell.

In approaching the eggshell as a material, we identify four main areas of investigation: (1) nucleation and growth of calcite from the membrane surrounding the egg white; (2) organic ultrastructure of the shell on which the mineralized layer grows; (3) shell porosity, including closed and open pores; and (4) the eggshell cuticle.

It is considered that, in living systems (3), biomineralization occurs from nanometric entities that produce hierarchical systems with interaction and feedback across different length scales. This approach has been suggested in studies on the eggshell and differs from composite technology (4,5). Decades ago (6), it was observed that both the organic component nucleates and grows on the inorganic substrate and that the organic phase may also nucleate on the calcitic substrate. This differs from composite technology, where at least two distinct entities are present: one phase (particle or fiber) and a matrix phase. In composite technology, wetting of the fiber by the matrix is essential. In eggshells of birds, chelonians, reptiles, etc., proteins appear to interfere with the thermodynamic phase (7–9), with the size and morphology of the polycrystals (10), and especially with the porosity of the inorganic matrix.

The classification of the eggshell as a composite appears in many studies describing biomineralization through heterogeneous nucleation of amorphous calcium carbonate (ACC) on the organic ultrastructure and its subsequent transformation into calcite. ACC would reach the biomineralization sites (membrane, palisade layer) in vesicles (11), with different proteins associated with the vesicle surface (12). As shell thickness increases, successive calcite layers nucleated and grown from the inner eggshell membrane form structures called mammillae and, micrometers above, the palisade layer develops (see Figure 9 in Results). In SEM images, a calcite crown (13) can be observed at the top of a mammilla (see Figure S7 in Supplementary Material). Although it represents less than 3% of the shell mass, a protein ultrastructure is associated with calcite nucleation and growth (10). How this occurs (sequentially or concomitantly) remains unclear.

One source of calcium (Ca) for egg production in laying hens (*Gallus gallus domesticus*) is the dissolution of medullary bone (14–16) and its transport via the bloodstream to the uterus, where biomineralization takes place.

The dissolution process of carbonated hydroxyapatite (CHAp) (17) from medullary bone and its transport through the bloodstream remain under investigation. Several published images show spherical hydroxyapatite na (13,18) nanoparticles on the inner side of the eggshell cuticle. Questions remain regarding the composition of these nanospheres, particularly whether phosphate occurs in inorganic and/or organic compounds and whether the nanospheres are crystalline or amorphous CHAp phases. Published infrared spectroscopy results (19) indicate that the nanospheres may be amorphous. However, in the pioneering work of Dennis et al. (13), images of hydroxyapatite nanoparticles are presented together with confirmation of hydroxyapatite crystallinity by X-ray diffraction.

More recent studies emphasize the initiation of nucleation of amorphous and/or crystalline calcium carbonate phases on the inner eggshell membrane, which contains the egg white and yolk (11,20–22). Little attention is given to the cuticle in these studies. Investigations focused on the cuticle are mainly concerned with effects of hen age, variations among breeds, and the impact of industrial egg-cleaning processes on the cuticle (19,23).

There is consensus regarding the presence of phosphate since the earliest studies (24) on cuticle composition. At one point, attempts were made to relate high phosphate concentrations to defects in the biomineralization process and even to premature oviposition (25,26), but no conclusion has clarified the role of phosphate in these phenomena.

Calcium phosphate nanospheres may be vesicles containing a liquid phase with aggregates of amorphous hydroxyapatite, similar to models proposed for bone biomineralization (27,28). Deformable nanometric vesicles facilitate transport through the bloodstream (especially in the microcirculation) and migration into the hen uterus.

Some recent studies (29) follow Dennis’ suggestion (13) that hydroxyapatite nanospheres in the cuticle would be involved in completing the calcite biomineralization process. Support for this hypothesis is indirect and derives from the observation (30) that phosphate would inhibit calcite nucleation.

Other recent studies (31) indicate that phosphate does not inhibit carbonate crystallization and, contrary to (30), may enable calcite crystallization in processes involving calcite formation through aggregation of amorphous nanoparticles.

The fact that calcium phosphate nanospheres are found in the cuticles of eggs from different bird species raised our initial questions about the biomineralization process in the avian uterus. Are these vesicles containing both mineral and liquid phases?

Is there a boundary between the cuticle (C) and the Vertical Layer (VL) that evidences a cuticle function of halting calcite growth? The suggestion that nanospheres serve to provide phosphate to inhibit calcite growth appears weakened in light of recent advances in the investigation of so-called non-classical calcite crystallization (31) in the presence of phosphates. In natural environments, phosphate presence leads to heterogeneous calcite crystallization (32).

It is also pertinent to investigate whether the calcium phosphate in the cuticle is B-type carbonated hydroxyapatite (CHAp-B), the mineral phase present in vertebrate bones, where approximately 6% of the PO₄³⁻ group is substituted by CO₃²⁻.

Studies of eggshell microstructure and nanostructure advocate the existence of hierarchical organization in shell porosity. Zhou et al. (33) studied pores that occur at high concentration and are distributed throughout the shell; their shape resembles bubbles and they exhibit similar submicrometric diameters. Studies on eggshell porosity are of both industrial and scientific interest (34–36). Using micro-CT, Riley et al. (37) found that void volume corresponds to approximately one-third of total shell volume. These voids are distributed throughout the shell. The images clearly demonstrate the gas-conducting role of open pores that traverse the entire shell and form through voids created between structures called mammillae. A thicker region of the cuticle is termed the pore plug (see Figure 3 in Results) and functions as a “cap” for this pore. AFM (Atomic Force Microscopy) has also been used to evaluate eggshell porosity (38).

Thus, the objective of this work was to conduct an extensive and detailed investigation by scanning electron microscopy (SEM), together with EDS, Raman, FTIR, XRD, and thermogravimetric analysis (TGA), to investigate the eggshell cuticle and, concomitantly, to attempt to elucidate:

i. the type of phosphate compound present in the cuticle;
ii. the interaction between the micrometric calcium phosphate layer and calcite;
iii. the connection between the inner calcium phosphate layer and calcite layer growth. During the course of the work, additional objectives were incorporated:
iv. to understand the growth morphology (uni-, bi-, or tridimensional) of calcite in the Vertical Layer (VL) and Palisade Layer (PL);
v. to understand VL and PL growth by addition of nanodroplets.

## 2. MATERIALS AND METHODS

### 2.1. Specimen Collection and Preparation

#### 2.1.1 Eggshells

Egg samples were obtained by purchase in supermarkets (white and brown eggs). The shells were washed under running tap water for a few minutes, followed by visual inspection to confirm the absence of any material adhered to the external and internal shell surfaces. Subsequently, the shells were dried in an electric oven (Brastemp, São Paulo, Brazil) at 60 °C for 3 hours.

After drying, the shells were stored in plastic bags and kept for SEM-EDS, Raman, FTIR, and XRD analyses. No additional thermal or chemical treatment was performed on the samples at this stage, which constituted the main part of the study.

For another stage of the study, eggshells were calcined in air at temperatures of 150, 350, 550, 750, and 950 °C in an electric furnace (Oga, Belo Horizonte, Brazil). After thermal treatment, part of the samples, cooled to room temperature, were ground in a porcelain mortar (Chiarotti, São Paulo, Brazil), while another part was kept as small fragments as removed from the furnace, which were very fragile. Subsequently, the samples (powder or fragments) were placed in Falcon-type tubes (15 mL) with caps and stored for subsequent TG, FTIR, and XRD analyses.

#### 2.1.2 Micro-Raman and Micro-FTIR Tests

For micro-Raman and micro-FTIR spectroscopy, eggshell samples were embedded in acrylic resin, sanded, and polished so that the shell thickness was exposed for analysis under the optical microscope of the instruments. Another type of mounting included fixing only part of the shell and performing very gentle sanding (followed by cleaning with extra-soft bristle brushes). This second type of preparation is shown in Figure 11.a.

#### 2.1.3 Chicken Bones

Chicken thighs were purchased in supermarkets and completely defleshed. The femur bones were brushed (plastic bristles) and washed under running water for a few minutes to remove adhered material. After this step, the bones were dried in an electric oven (Brastemp, São Paulo, Brazil) at 60 °C for 3 hours.

After drying, the bone samples were immersed in a 2.5% w/w sodium hypochlorite solution for 72 hours. The samples were then dried again at 60 °C for 3 hours. The pieces were broken, and the fragments were placed in Falcon tubes and kept under refrigeration until Raman and FTIR spectroscopy analysis.

#### 2.1.4 Bovine Hydroxyapatite

Bovine hydroxyapatite was obtained in the laboratory using bovine tibia pieces supplied by a butcher, already cut into cylinders 3–4 cm in length. Initially, they were cleaned with a brush and running water and left for 6 hours at 60 °C in an electric oven (Brastemp, São Paulo, Brazil) to remove viscous components from inside the tibia. After this step and another brushing under running water, the pieces were dried again at 60 °C for 3 hours in the same oven.

After drying, the bone samples were immersed in a 2.5% w/w sodium hypochlorite solution for 96 hours. The samples were then dried again at 60 °C for 6 hours. The pieces were broken, and the fragments were stored in Falcon tubes under refrigeration. For X-ray diffraction analysis (Supplementary Material, Figure S8), the fragments were pulverized.

### 2.2. X-Ray Diffraction (XRD)

For X-ray diffraction analysis of the crystalline phases present in eggshell samples, a multipurpose diffractometer (Anton Paar, model XRDDynamic 500, Graz, Austria) equipped with a copper tube was used. Experimental parameters were 40 kV voltage, 50 mA cathode current, and a 2θ angle range of 10–90°. Phase identification was performed using the Inorganic Crystal Structure Database (ICSD), accessed through the CCDC software platform. For the tests, the eggshells were again carefully washed with deionized water and a brush and air-dried. Analyses were performed on eggshell powder ground in a mortar and pestle as described above (item 2.1).

### 2.3 Scanning Electron Microscopy (SEM) with Energy Dispersive Spectroscopy (EDS)

Preparation and analysis of eggshell samples by SEM were conducted at the Microscopy Center of the Federal University of Minas Gerais (CM-UFMG) using three instruments:

a. FEI Quanta 3D FEG dual-beam (Hillsboro, USA) with backscattered electron (BSE) and EDS detectors;
b. APREO-2C ThermoFisher (Waltham, USA) with standard BSE and EDS detectors;
c. FEI Quanta 200 FEG (Hillsboro, USA).

Three eggshell specimens were washed under running tap water for a few minutes and then visually inspected to confirm the absence of any material on the external and internal shell surfaces. The shells were subsequently dried in an electric oven at 60 °C for 3 h. After drying, the samples were coated with a 50 nm graphite layer. A fourth specimen, prepared under the same conditions, was coated with gold–palladium. All samples were stored in a low-humidity desiccator before and after the microscopy sessions. Six sessions of 2 hours each were carried out to investigate only the eggshell thickness. Elemental maps of carbon (C), oxygen (O), calcium (Ca), phosphorus (P), magnesium (Mg) and sulphur (S) show slight color differences among figures due to SEM-EDS sessions conducted on different dates.

### 2.4 Raman Spectroscopy

Raman spectroscopy analyses were performed using two instruments:

*2.4.1*- Spectrometer Horiba Jobin Yvon (Oberursel, Germany) LabRam-HR 800 equipped with a He-Ne laser (632.8 nm excitation) and an Olympus BX41 microscope with 10×, 50×, 100×, and 100× LWD objectives. The laser was focused on an area of 1–2 µm² (100× objective). Scattered light was collected by a monochromator and detected by a liquid nitrogen-cooled CCD. Spectra were recorded from 100 to 4200 cm⁻¹ with 1.1 cm⁻¹ pitch. Most measurements were acquired in the 80–1250 cm⁻¹ range, with 30 s acquisition time and 10 accumulations to improve signal-to-noise ratio.

*2.4.2*-Witec (Ulm, Germany) Alpha300 RA spectrometer with a 532 nm diode laser coupled to an optical microscope and EMCCD detector. The beam was focused on approximately 1 µm². Spectra were mainly acquired between 80–1250 cm⁻¹, 1 cm⁻¹ resolution, 1800 lines/mm grating, 100× objective, 10 accumulations, and 60 s integration time per accumulation.

In both instruments, laser power was adjusted to avoid local heating of the samples.

### 2.5. FTIR and Micro-FTIR-ATR Spectroscopy

FTIR and FTIR-ATR analyses were performed on eggshell samples prepared as described in item 2.1, either pulverized or in fragments approximately 2 cm in edge length. Untreated samples and samples thermally treated at 150, 350, 550, 750, 950°C were analyzed. Treated femur fragments and bovine hydroxyapatite samples were also analyzed.

Infrared spectroscopy was performed using:

*2.5.1.* IRSpirit FTIR (Shimadzu, Kyoto, Japan), 400–4000 cm⁻¹ range, transmittance mode (%);

*2.5.2.* Tracer-100 FTIR-ATR (Shimadzu, Kyoto, Japan) coupled to AIM-9000 (Shimadzu, Kyoto, Japan) infrared microscope with reflectance and ATR modules, 700–4000 cm⁻¹ range, transmittance mode (%). All measurements were conducted at room temperature. Powder spectra were recorded at 2 cm⁻¹ resolution; FTIR-ATR spectra at 1 cm⁻¹ resolution. Spectral interpretation was performed using the Cambridge Crystallographic Data Centre (CCDC) software. FTIR results are presented in Section 3.6 (Results) and in Section 3 of the Supplementary Material.

### 2.6 Thermal Analysis (TGA)

Thermal analysis was performed using a TG/DTA Shimadzu (Kyoto, Japan) model DTG-60/DTG-60H. An alumina crucible (Al₂O₃) was used under nitrogen atmosphere (100 mL min⁻¹) with a heating rate of 10 °C min⁻¹. The temperature range investigated was from room temperature to 600 °C. TG results are presented in the Supplementary Material.

### 2.7 Software for Measurements (Feret Diameter) and Statistical Analysis

Image measurements from SEM images were performed using ImageJ2 version 2.16.0/1.54p. Statistical analysis was carried out using StatPlus v. 8.0.4.0. The Kruskal–Wallis significance test followed by Dunn’s post-test with Bonferroni correction was applied to analyze the dimensions of structures and morphologies present in the investigated eggshell samples.

## 3. RESULTS

### 3.1. SEM–EDS

Figures 1 to 9 present the results of the SEM image analysis and the EDS spectroscopy coupled to SEM. We intentionally captured images of the same scene using two types of detectors: the secondary electron detector (SE), which details the sample topography, and the backscattered electron detector (BSE), which converts compositional differences into light/dark contrast, enabling image contrasts that support a detailed understanding of the scene. What appears to be a solid surface in SE is revealed, in the BSE sensor, as much more detailed surfaces and subsurfaces.

**Figure 1.**
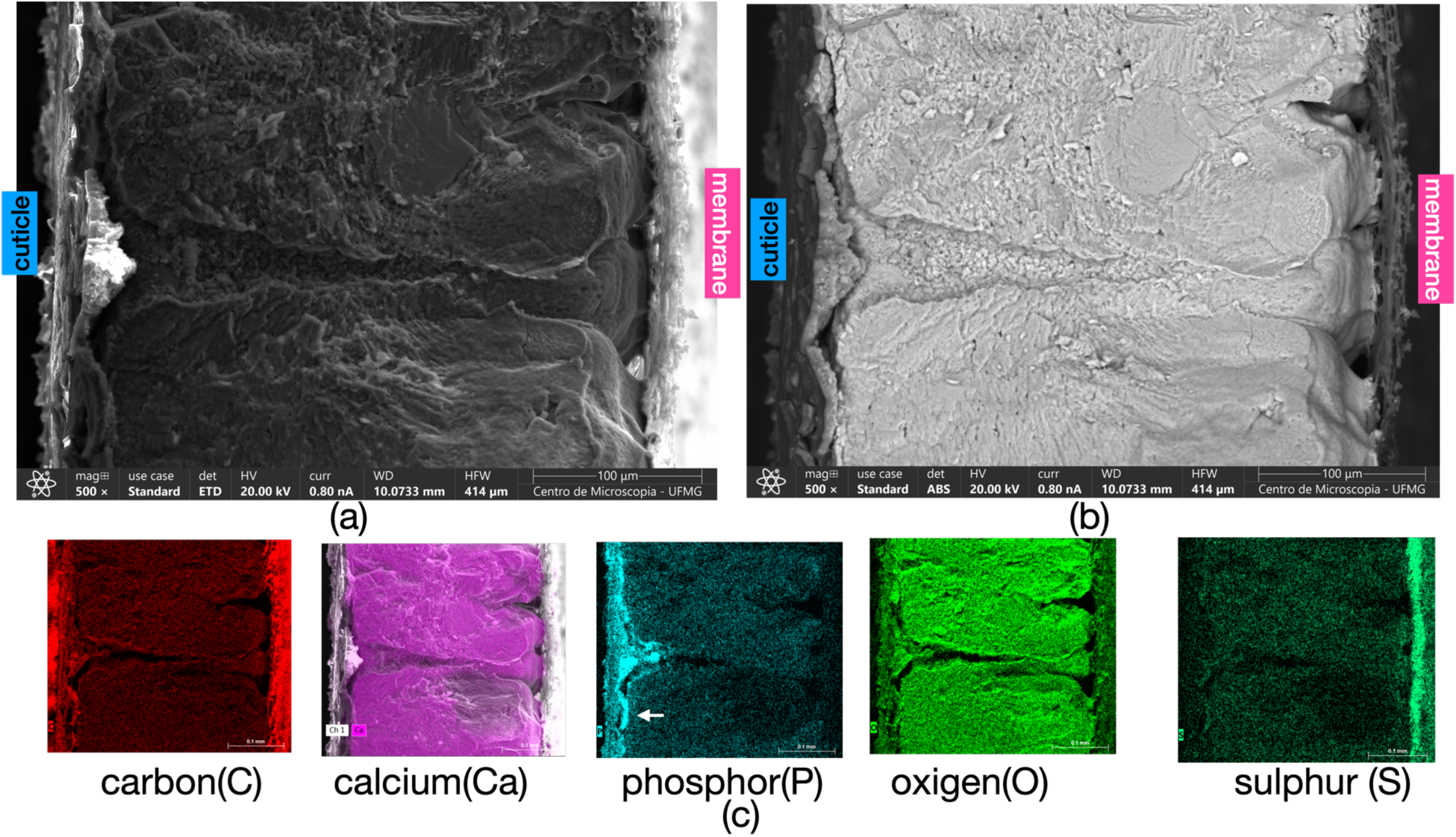
(a) Image of the thickness [(353 ± 7) μm] of a laying hen eggshell (*Gallus gallus domesticus*), from the cuticle (left side) to the membrane (right side) where biomineralization begins. The SE (secondary electrons) detector highlights the shell topography. It is possible to observe, in the central part of the image, the presence of voids forming channels/pores that extend uninterruptedly from the cuticle to the membrane.Image (b) shows the same scene of the same sample, but using the BSE (backscattered electrons) detector, with lighter regions corresponding to chemical elements of higher atomic weight. It is possible to observe (left side of the image) that the tips of the mammillae begin at different points adhered to the membrane surface.(c) Profiles of the most abundant chemical elements in the eggshell: C (carbon), Ca (calcium), P (phosphorus), O (oxygen), and S (sulfur). While carbon (C) and oxygen (O) are found on both faces of the shell (upper part of the cuticle and inner membrane), phosphorus (P) is concentrated in the cuticle and calcium (Ca) throughout the entire biomineralized extension. The white arrow in the phosphorus (P) profile indicates a higher concentration of this element on the lower face of the cuticle. Observe, in the same image, that the phosphorus-containing layer (pore plug) presents a larger area in the region of the channel/pore that crosses the shell.

In Figure 1 we present images of the laying-hen eggshell throughout its entire extent, from the cuticle to the inner membrane. The total thickness of this eggshell was evaluated as (353 ± 7) μm (see Table 1), including: a—cuticle (C), b—vertical layer (VL), c—palisade layer (PL), d—mammillary region (MA), and e—membrane (M). In the center of the image, channels crossing the entire eggshell thickness can be seen. These channels have substantial widths (∼70 μm) both near the inner membrane and near the cuticle. Their three-dimensional shape would be like two cones whose apices meet close to mid-thickness. The structures called mammillae, where calcite (CaCO₃) nucleation and growth occur, are clearly visible in both images of Figure 1. In 1b it is possible to observe that the bases of the so-called mammillae (39) start at individualized and distinct points adhered to the external face of the inner membrane. According to the literature, the inner eggshell membrane (40) is in fact composed of two membranes, one inner and one outer, both made of collagen (41).

**Table 1.**
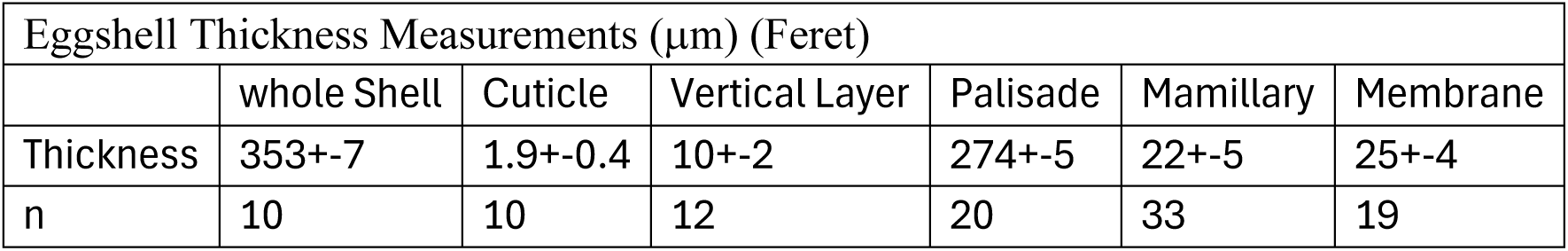
Measurements of eggshell thickness and its layers using SEM images and ImageJ software.

Elemental maps of carbon (C), calcium (Ca), phosphorus (P), oxygen (O), and sulfur (S) in the eggshell, obtained by energy-dispersive spectroscopy (EDS), are shown in Figure 1c. A white arrow indicates the location of phosphorus (P) in the lower part of the cuticle. In the channels that cross the shell, the thickness of the phosphorus-containing layer, also called the pore plug (42), extends further into the shell interior. Carbon (C) appears marked on the upper face of the cuticle and on the inner membrane, whereas calcium (Ca) and oxygen appear marked throughout the shell thickness. Also note the presence of sulfur (S)—due to sulfated polysaccharides (6)—in the inner eggshell membrane.

Table 2 presents the concentration data (EDS, semi-quantitative) for carbon (C), oxygen (O), magnesium (Mg), phosphorus (P), sulfur (S), and calcium (Ca). The data are divided by the investigated areas (cuticle, cuticle + vertical layer VL, etc.; see the legend on the right side of Table 2).

**Table 2.**
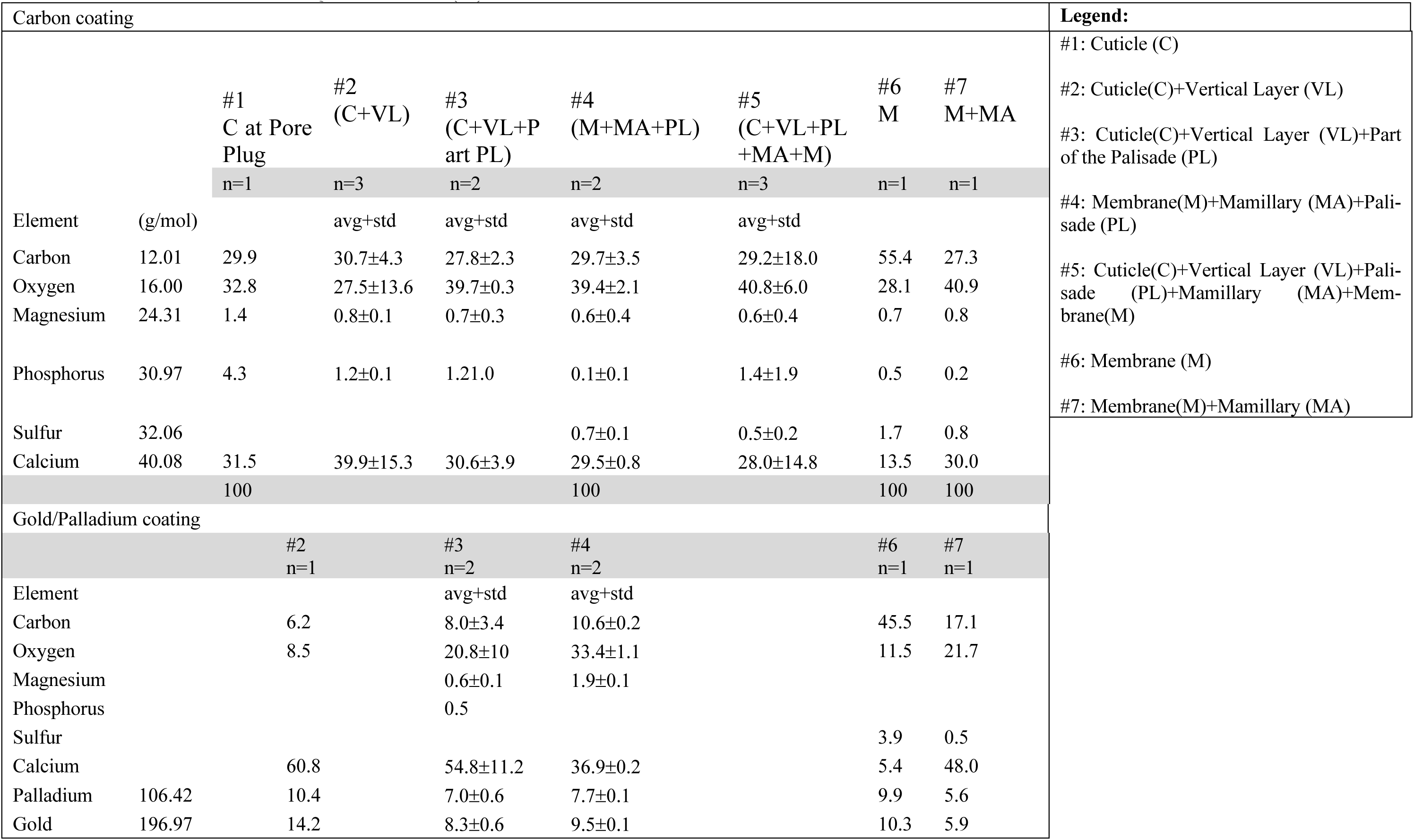
EDS-Elemental Mass Quantification (%)

In Table 2, the two types of sample coating (carbon or gold/palladium) are separated. Carbon coating is the preferred option for EDS because it does not generate interfering peaks, allowing more accurate quantification. On the other hand, gold peaks overlap with those of other elements such as phosphorus. In EDS measurements performed with gold coating (lower part of Table 1), phosphorus is practically absent because the gold peak overlaps it. Figure S6 (Supplementary Material) shows the overlap of the gold peak with phosphorus in the EDS analysis of the cuticle. This may be one source of the non-detection of phosphorus in vesicles that would transport material for biomineralization (43).

The SEM images in Figure 2 correspond to the cuticle (black arrow in a1) and to the calcite region immediately below the cuticle (marked by magenta ellipses in a1 and b1). The cuticle thickness was evaluated as (1.9 ± 0.4) μm. Images from left to right correspond to progressively higher magnifications of the same region, focusing on the cuticle (magenta arrows). Cuticle thickness is shown in a2 and b3 with black arrows and in a3 with white arrows. Images a1, a2, and a3 were acquired with the SE sensor (topography), whereas images b1, b2, and b3 were acquired with the BSE sensor, which allows compositional differences to be distinguished as darker or lighter regions depending on the atomic weight of the elements present. Thus, the calcite layer appears light gray, whereas the cuticle appears dark gray. In images b1, b2, and especially b3, spheres of different diameters are observed throughout the cuticle (see the plot with the diameter of all spheres and semi-spheres found across the shell thickness in Table 3 and Figure 10).

**Figure 2.**
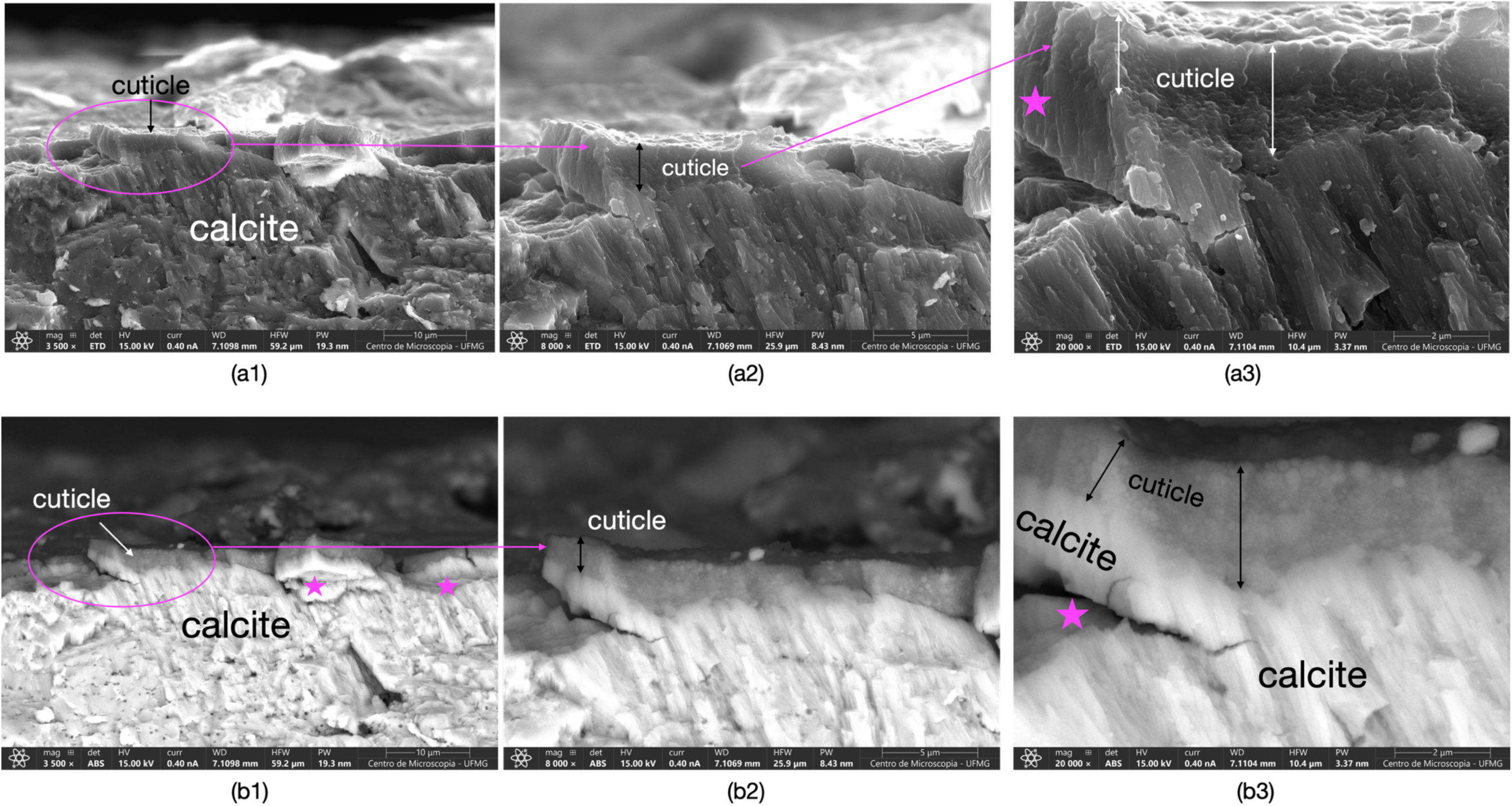
SEM images, at different magnifications, of the same area of the eggshell cuticle: SE detector images (a1, a2, and a3) providing the topography of the eggshell cuticle, and images b1, b2, and b3 acquired with the BSE detector, providing more detailed images (compositional differences) of the same region. The presence of spheres in the cuticle is visible especially in images b2 and b3. Spherical shapes in the cuticle region can also be observed in image a3. A magenta asterisk was placed at locations where fractures evidence continuity between the cuticle and the underlying calcite vertical layer (VL).

**Table 3.** Mean and standard deviation of the Feret diameter (nm) of spherical and hemispherical struc-tures measured in the different layers of the eggshell; measurements of the thickness of an individual lamella (nm), which constitutes the palisade, are presented in the last row of the table.

| Feret diameter (nm) |  |  |  |  |  |
| --- | --- | --- | --- | --- | --- |
|  | eggshell layer | shape | mean | std | count |
| item |  |  |  |  |  |
| 1 | Cuticle | spheres | 96 | 52 | 84 |
| 2 | Vertical Layer | hemispheres | 88 | 29 | 52 |
| 3 | Palisade Layer | hemispheres | 57 | 12 | 35 |
| 4 | Pore Plug | spheres | 254 | 110 | 18 |
|  |  |  | thickness(nm) |  |  |
| 5 | Palisade Lamella | lamella | 30 | 12 | 38 |

A magenta asterisk was placed in a3 and b3 to show that a fracture in the cuticle region kept the cuticle and the vertical calcite layer (VL) united and intact, while the fracture propagated through the VL. In image b3, acquired with the BSE sensor, unidimensional (fibrillar) calcite growth is observed, intimately connected with the base of the cuticle.

In Figure 3, the images show the cuticle in the region of the channels presented in Figure 1. These channels, called pores in the literature (40,42), occur as a consequence of the mammillae shape: they are wider both in the mammillary region and in the upper part, just below the cuticle. Images a1 (SE sensor, topography) and a3 (BSE sensor, greater depth in structural detailing) show that the cuticle has a much greater thickness at this location, forming a “cluster” of spheres. The EDS composition maps shown in a2 and b2 confirm phosphorus (P) as one of the elements present in the spheres. Images b1 (SE sensor) and b3 (BSE sensor) show in detail the spheres, with similar diameter, that form this “cluster”.

**Figure 3.**
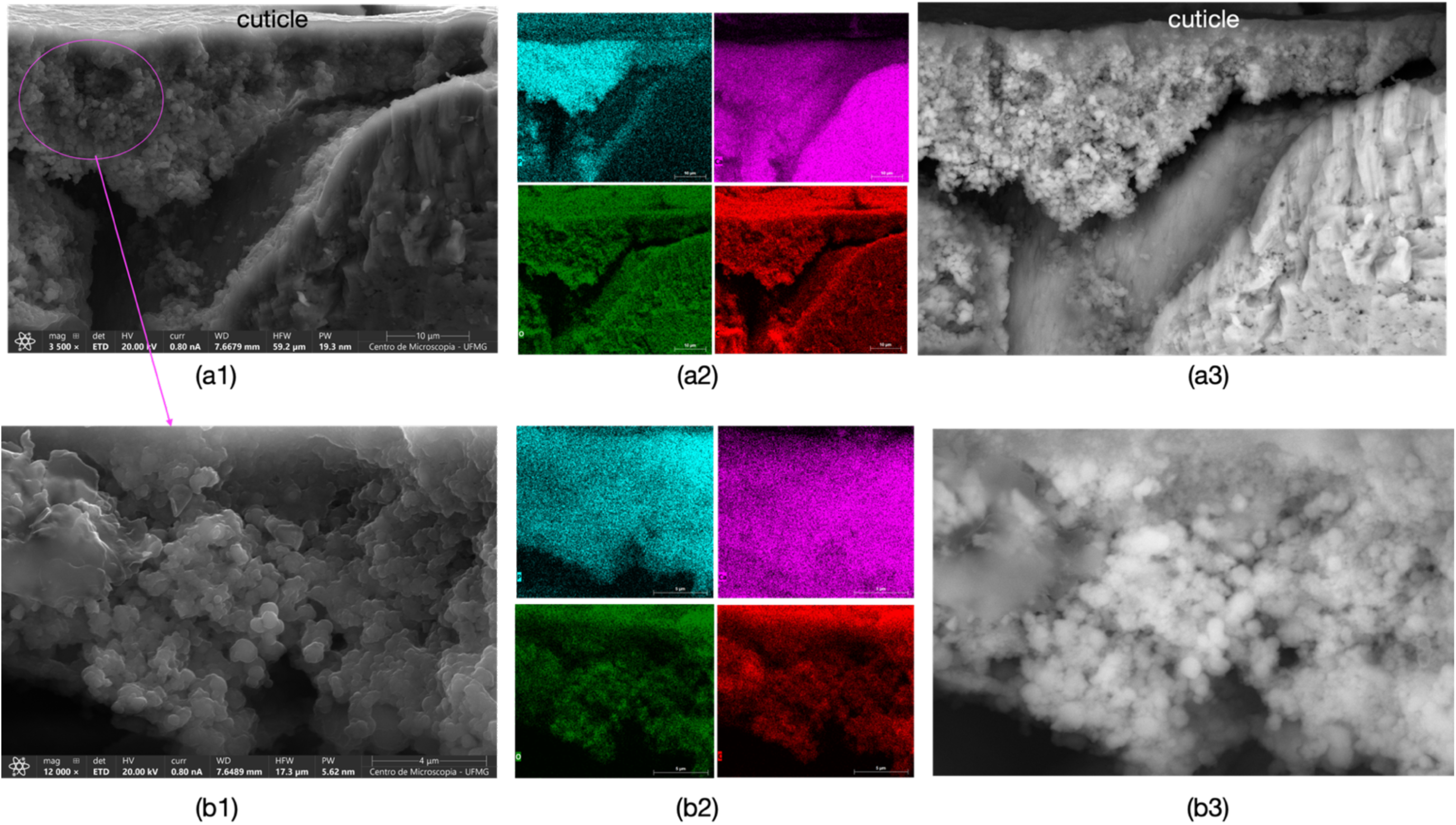
SEM images of the cuticle located exactly above a channel/pore that crosses the entire shell thickness (region of the “pore plug” (42)): (a1) SE detector; (a2) EDS spectroscopy map showing the distribution of phosphorus (P, light blue), calcium (Ca, magenta), oxygen (O, green), and carbon (C, red); (a3) same area as (a1) acquired with the backscattered electron (BSE) detector. (b1) shows a detail of (a1) (region circled in magenta). Spheres are visible in the images; (b2) EDS map showing the distribution of P, Ca, O, and C in the area presented in image (b1); (b3) BSE detector: the presence of spheres in the cuticle is clearly visible. The semi-quantitative (EDS) concentration of the elements P, Ca, O, and C present in these areas is presented in Table 2, and the diameter of these submicrometric spheres is shown in the graph of Figure 10. In (a1) and (a3), the spheres are arranged as a “cluster of grapes.” This “cluster” occurs, as observed above, in the region of channels extending from the inner membrane to the cuticle (shown in the center of the whole-shell images in Figure 1).

The images in Figure 4 show the cuticle and the cuticle (C) together with the vertical layer (VL), with 4a showing in detail the spheres in this layer. The spheres on the eggshell surface have dimensions close to 300 nm and would have, among other functions, protection against solar radiation and infectious agents (25,43,44). Also in 4a it is observed that smaller spheres are located throughout the cuticle but predominate on the lower face. The diameter of these smaller spheres was evaluated as (96 ± 52) nm and is shown in Table 3 and Figure 10. Image 4b (BSE sensor) shows both the cuticle region and part of the VL, whose thickness was evaluated as (10 ± 2) μm. The fibrillar aspect of the VL and its interaction with the cuticle (C) nanospheres can be observed. Images 4c1 and 4c2 are, respectively, SE (topography) and BSE images showing both the VL and part of the palisade layer (PL). The PL presents plates, pores, and different growth directions of the calcitic layer, whereas the VL presents essentially a single direction, possibly due to its thickness of only about a few micrometers

**Figure 4.**
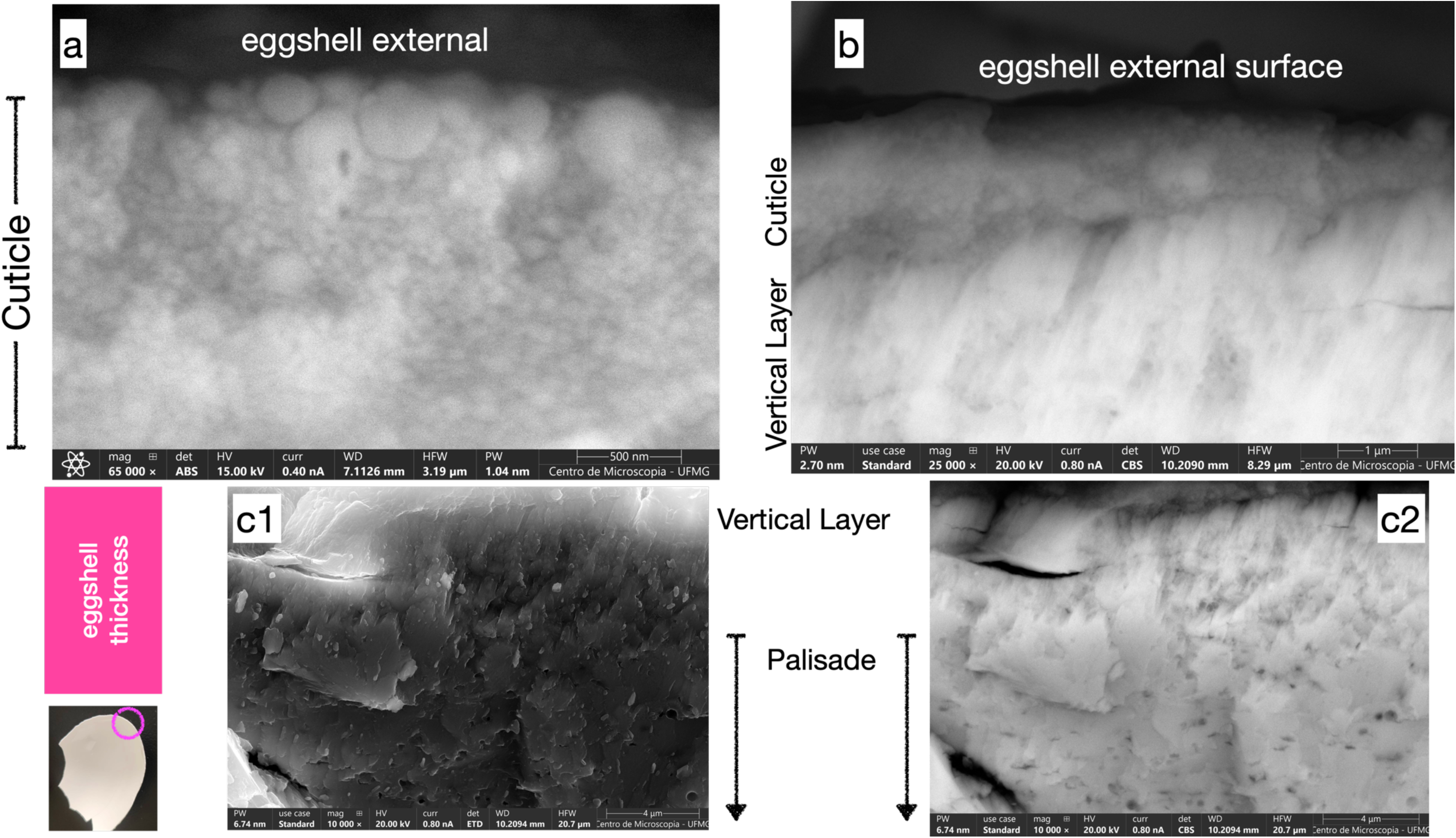
SEM images of the shell thickness: (a) Cuticle surface (BSE) showing larger spheres (∼300 nm) on the surface. The diameters of the smaller spheres, nanospheres, were evaluated as (65 ± 24) nm (see Figure 6). (b) Cuticle (thickness = 1.9 ± 0.4 μm) and part of the vertical layer (VL). The fibrillar aspect of the vertical layer is visible in the BSE image. (c1) Vertical layer VL (thickness = 10 ± 2 μm) and part of the palisade layer (PL). (c2) Same image, BSE detector. The palisade (lower part of the image) appears compact both in the surface (SE) image and in the BSE image.

#### 3.1.2. Hemispheres (nanodroplets before solidification) in the Vertical (VL) and Palisade (PL) Layers

Images 5a and 5b are topographic (SE), whereas 5b1 was acquired with the BSE sensor. A white circle encloses some nano-hemispheres (and which had previously existed as <u>nanodroplets</u> before solidifica-tion) adhered to the plates of the vertical layer (VL). The discovery of these morphologies occurred when we were investigating palisade pores. These nanodroplets appear to indicate the shell-thickening route, which occurs by deposition and coalescence of nanodroplets (visible inside and outside the white circles in 5a and 5b) in both the VL and the palisade PL (images 5c and 5d, topography). The diameters of these nanodroplets or nano-hemispheres (Table 3 and Figure 10) were evaluated as (88 ± 29) nm in the VL and (57 ± 12) nm in the PL. After coalescence, diameters are larger. Images 5b1 and 5d1 were acquired with the BSE sensor and correspond to the same scene (yellow bidirectional arrows), evidencing structures present in areas identical to those of images 5b and 5d. It is possible to observe, especially in b1, that the plates seen in image 5b are formed by unidimensional (fibrous) calcite structures that grow through mutual adhesion of these nano-hemispheres.

We call them nanodroplets because, in the liquid environment of the laying hen uterus, these nanospheres, upon adhering to the already solidified structure, assume a droplet-like shape attached to the substrate. There are nano-hemispheres even in the cuticle–vertical layer (VL) interaction region. These nano-hemispheres or nanodroplets must therefore contain both calcium carbonate (ACC?) and liquid, since they are deformable and coalesced on the pre-existing surface. In the already formed palisade PL, fiber density is much higher than in the VL, as can be seen by comparing Figure 5d1 with 5b1. In 5d1, for example, it is only possible to distinguish lower fiber density at the pore edges (upper part at the center of image 5d1).

**Figure 5.**
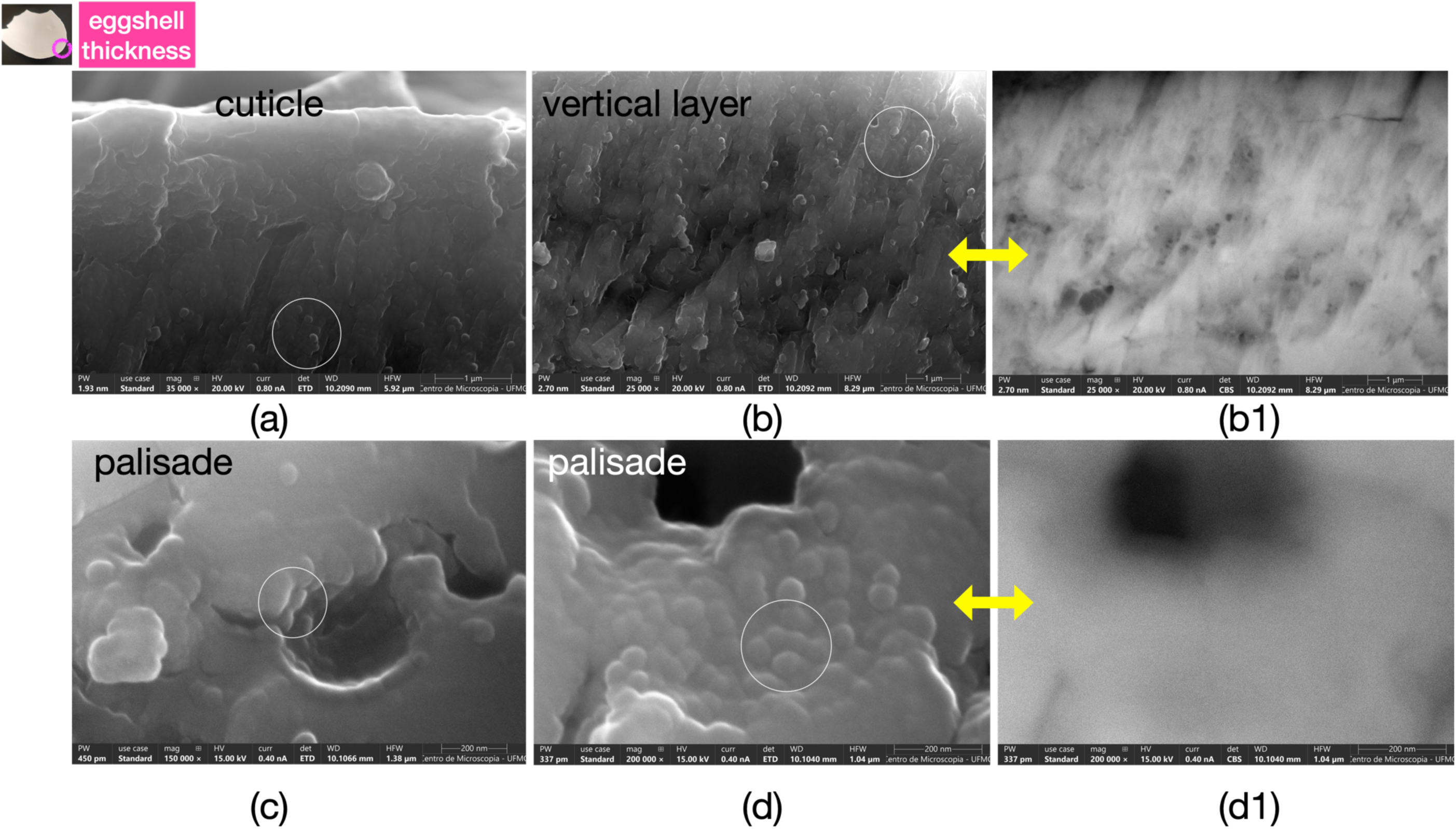
SEM images showing details of eggshell thickening growth occurring through deposition and coalescence of nano-hemispheres (marked with white circles), located both in the vertical layer (images a and b) and in the palisade layer (images c and d). Nano-hemispheres have diameters approximately between 50 and 90 nm. After coalescence, the diameters are larger. Images a, b, c, and d are topographic images acquired using the SE (secondary electrons) technique. Images b1 and d1 were acquired using the BSE (backscattered electrons) technique, which indicates compositional differences, and correspond to areas identical to those shown in images b and d. It is possible to observe, especially in b1, that the plates seen in image b contain unidimensional entities (fibers) of calcite, with fiber diameters on the order of a few nanometers. The nano-hemispheres must therefore contain both calcium carbonate (ACC?) and organic material (proteins and polysaccharides?). In the palisade PL, the density of calcium carbonate appears much higher than in the vertical layer, as can be concluded by comparing Figure d1 with b1.

Images 6a and 6b show the palisade layer (PL) (39,40). The total thickness of the PL was evaluated as (274 ± 5) μm (Table 1) and represents nearly 80% of total shell thickness. In the palisade PL, lamellae grow over one another and in different directions. For example, in image 6a the darker gray region (lower left) grows almost perpendicular to the growth of the lamellae present in the remainder of the image. A detail of this region is shown in 6b. The nanometric lamellae thickness, in the samples investigated in this work, was evaluated (see Table 3) as (30 ± 12) nm. The high standard deviation of these measurements results from the difficulty in individualizing lamella thickness. Pink asterisks in 6b highlight how thin (and nearly transparent) these nanometric structures are. Even in an SE topography image, which captures electrons from depths <10 nm, it is possible to visualize details below the lamella. In 6c (BSE image), this lamella transparency due to their minute thickness is more evident because of the greater depth captured by the BSE sensor. It was not possible to evaluate lamella width, and they appear to vary in density (or composition) along their width.

Figure 7 presents images of the palisade PL region showing growth of layers in different directions and senses (Figure 7a). Black points in Figure 6a correspond to pores already seen in Figures 5a, 5b, and 5c. Figure 7b evidences a large number of circular pores—closed or communicating with adjacent pores—present throughout the region where calcite fibrous growth occurs. From the images, it is not possible to observe any preferential porosity orientation. Comparing the pores inside the red circle in Figure 7c (SE, topography) with the same pores (red circle in Figure 7d) obtained with the BSE sensor, it is observed that the pores cross dozens of lamellae, have nearly spherical morphology, and represent—together with the channels shown in Figure 1—a considerable fraction of the shell volume (36,45,46).

**Figure 6.**
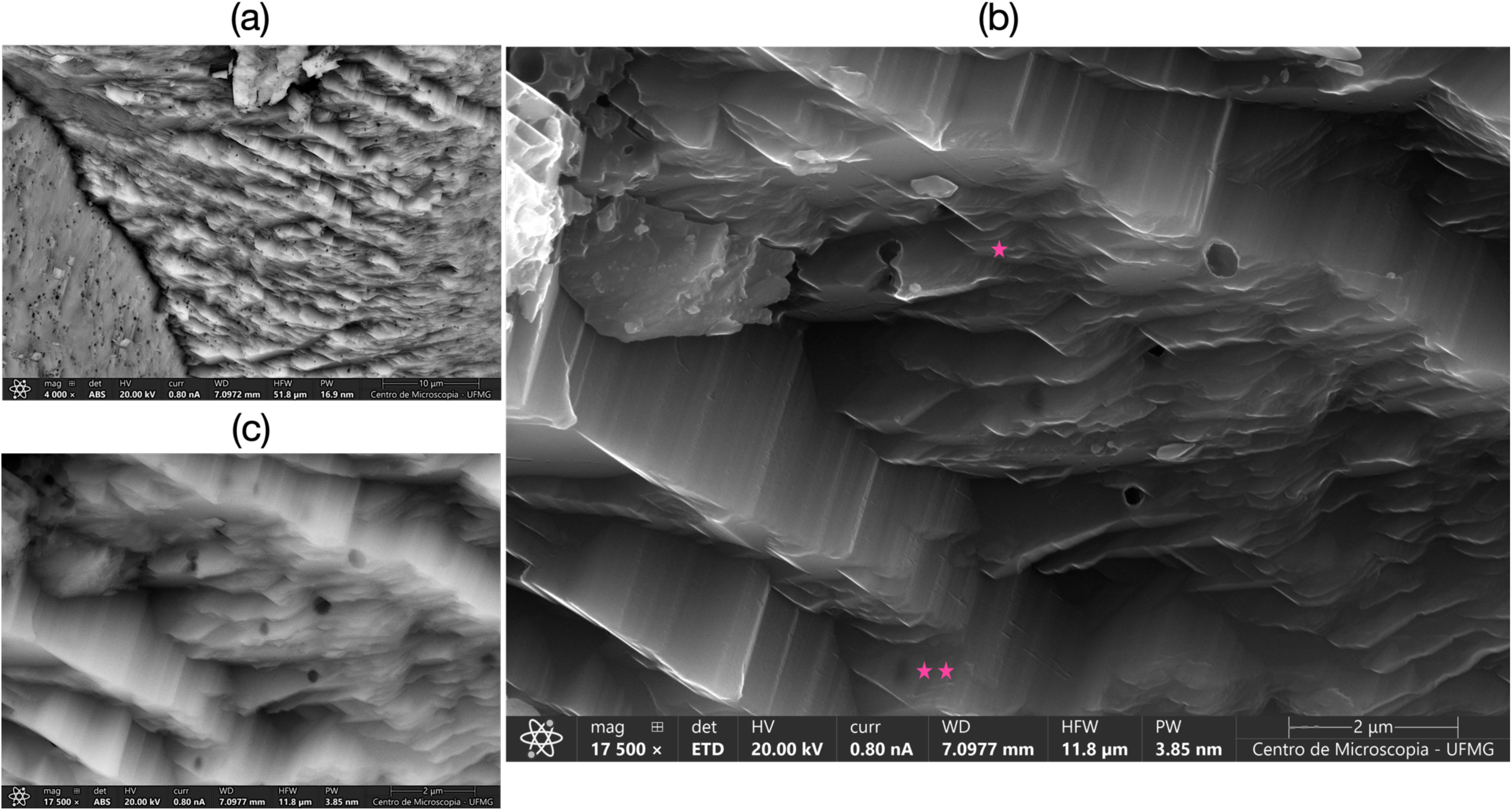
(a) Fracture image of the eggshell in the palisade layer (PL) region showing nanometric-thickness lamellae evaluated as 30 ± 12 nm. A large number (possibly several hundreds) of individualized and superposed lamellae are observed on the right side of the image. Note the enormous number of pores within the lamellae; (b) Detail of the superposed lamellae in the PL. Because they are so thin (pink *), some lamellae are transparent, revealing underlying pores (**) even in SE mode, which captures the electron beam at depths of 1–5 nm within the sample. The unidimensional and anisotropic growth, in fibers, as shown in Figures 4b and 5b1, possibly adhered to each other by surface interactions, water, and other substances, forms the lamellae observed in the images. Fiber growth appears to be consistent both with the speed of shell biomineralization and with its mechanical resistance; (c) Same image as (b), but acquired with the BSE detector. Lamella transparency is greater due to the deeper beam penetration captured by the BSE sensor.

**Figure 7.**
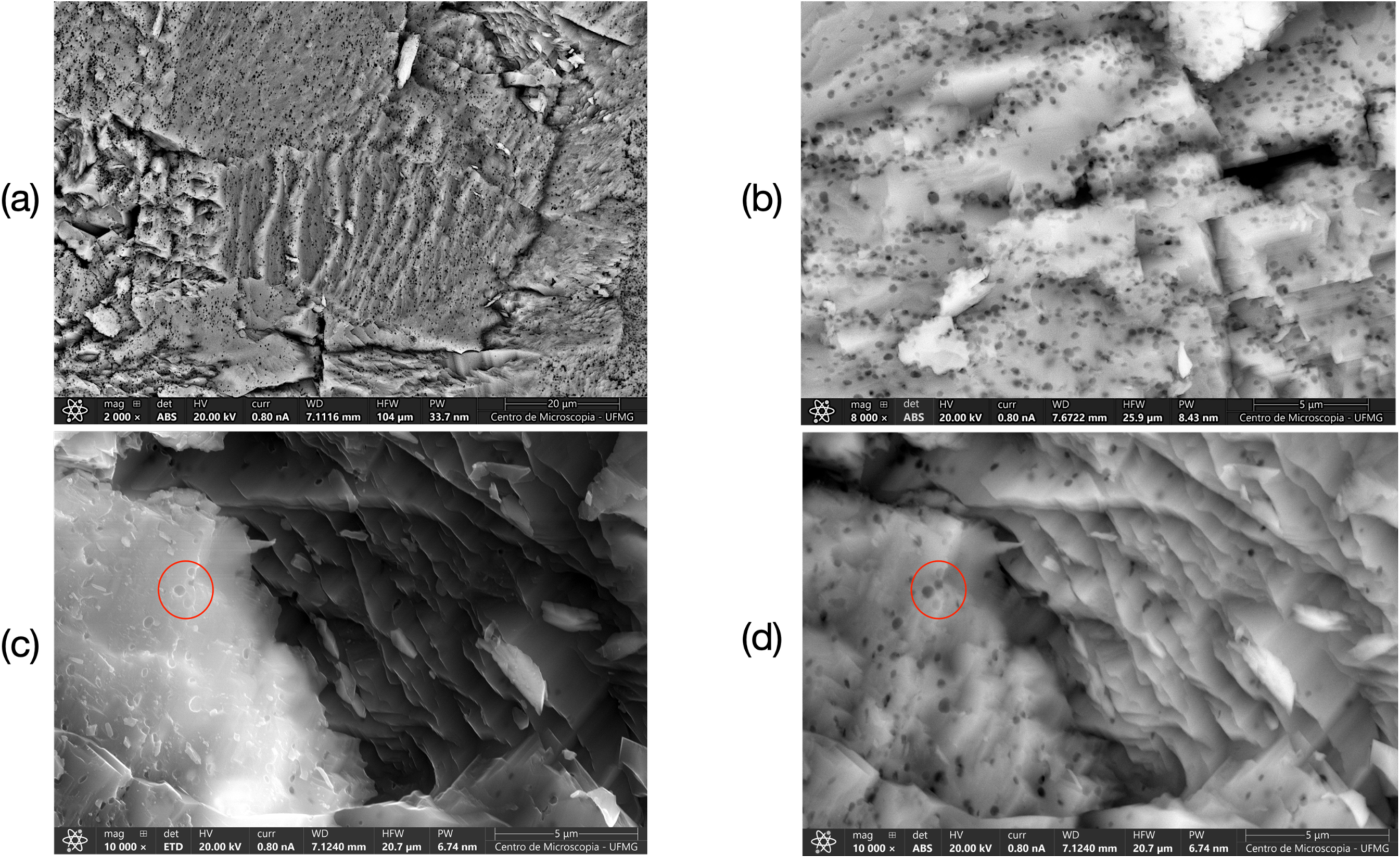
(a) Image, acquired using the BSE sensor, of a fractured shell area with the crystallized calcite region showing a significant number of circular pores; (b) High porosity of the calcite layer. An evaluation of porosity (P) presented in the two-dimensional image indicated 20% < P < 33% (Figure S8 in the Supplementary Material); (c) and (d) are images of the same calcite area, with (c) acquired using the SE sensor, highlighting topography, and (d) acquired using the BSE sensor with greater depth. The pores occur in the calcite growth plane. Within the red circle (left side of the figure), at least four pores are clearly visible. Note that the pores have approximately circular shapes. Two pores appear deeper (darker), as shown in (d), while two are shallower, indicating that they are spheres cut (by fracture) at different heights. Pores exist where previously there was a liquid or a gas. In this case, possibly liquid, because biomineralization occurred in the uterus, in an aqueous environment. However, calcite growth was not interrupted by the pores, as evidenced by the images in Figures 5 and 8 (below).

Figure 8a shows a palisade PL region containing several pores. The white numbering 1, 2, and 3 in image 8a indicates regions shown in detail in images 8b, 8c, and 8d (SE, topography). Images 8b′, 8c′, and 8d′ correspond to the same areas but acquired with the BSE sensor. The nearly spherical pores form concomitantly with the growth process of superposed lamellae. In 8c′ (BSE), lower right corner, it is possible to observe the presence of a pore in the lower left corner of the image that was later covered by superposed lamellae (image 8c). In 8d′, a void located at mid-height of the right border is not visible in 8d because it was covered by superposed lamellae. Pore edges in 8b, 8c, and 8d are filled with nano-hemispheres or nanodroplets that coalesced to form a new calcite lamella. In the PL, the individual diameters of these hemispheres were evaluated as (57 ± 12) nm (see Table 3 and the plot in Figure 10). We named them nanodroplets because they were viscous and deformable suspensions during the process of calcite lamella growth, which proceeds by additive growth through nanodroplet coalescence in the palisade layer PL and in the VL. In 8b, upper left corner (just below the letter b), lamella growth occurs by coalescence of these nano-hemispheres. In 8c, in the lower part of the image, nanodroplets are coalescing and forming superposed layers. In 8d, this lamella growth by coalescence of nanodroplets is also observed, with a lamella growing over incomplete underlying lamellae (located at mid-height of the right border of the image).

**Figure 8.**
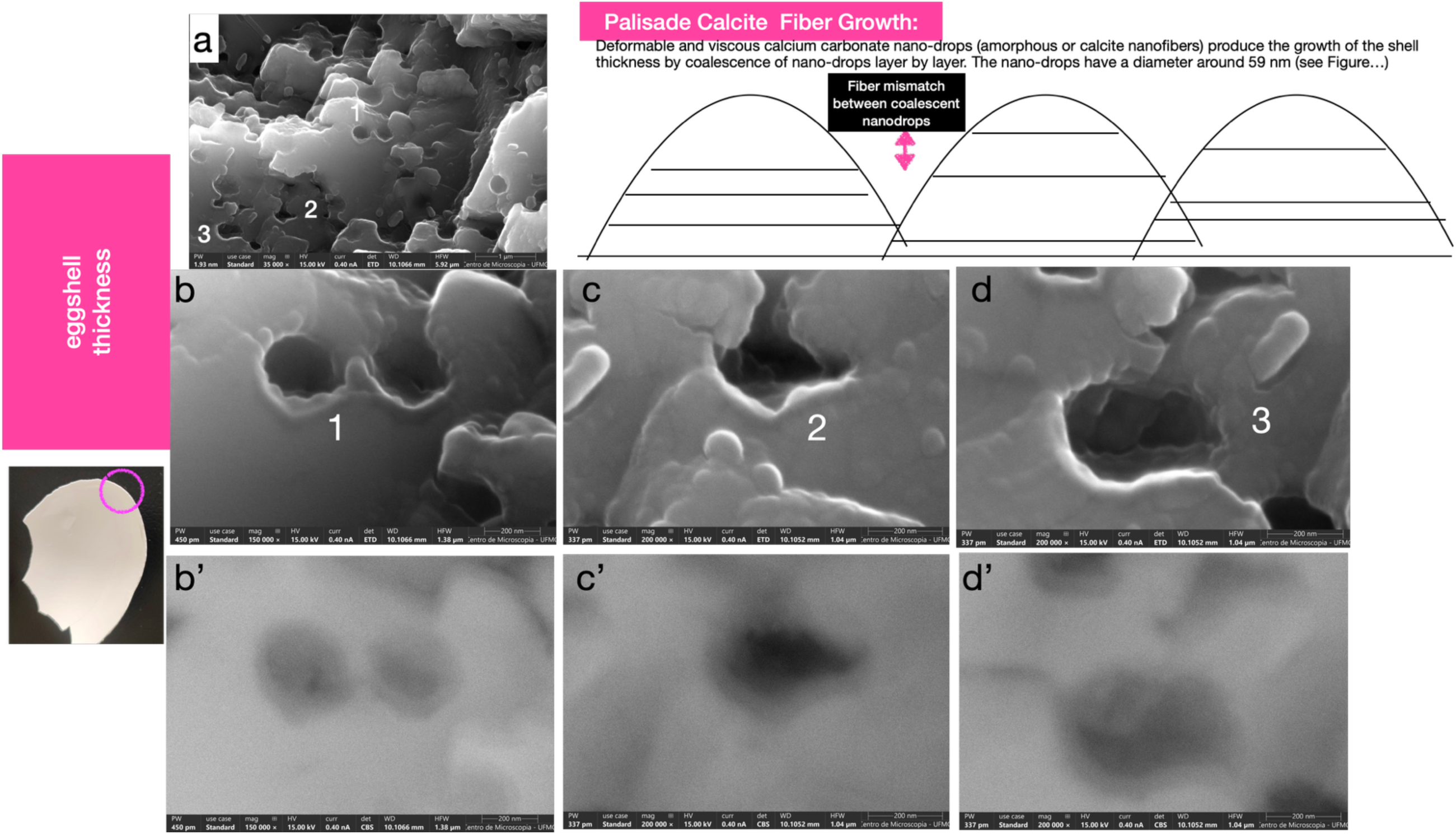
SEM images obtained across the shell thickness: (a) Region of the palisade layer, SE sensor, surface topography. The numbers 1, 2, and 3 represent areas shown in detail in images b, c, and d, also acquired using the SE sensor. Images b′, c′, and d′ correspond, respectively, to the same areas as b, c, and d, but obtained using the backscattered electron (BSE) detector. By comparing the SE and BSE images, it is possible to observe that the nanodroplets are suspensions, since they are deformable and coalesce, as shown in image b at the pore edge near number 1, in image c in the cluster of nano-hemispheres near number 2, and in the coalescence of nano-hemispheres at the pore edge shown in image d, near number 3. The BSE images shown in b′, c′, and d′ indicate (especially within the pores) unidimensional growth of calcite, in fibers, as observed in the vertical layer (Figures 4b, 5b1, and 6d). The presence of nano-hemispheres after biomineralization was completed indicates that there was not enough time for them to fully coalesce into a nanolayer of calcite. It may also indicate that the limiting factor is misalignment between fibers existing within each nano-hemisphere during coalescence, as suggested in the drawing (upper right part of the figure) and evidenced by the micrometric length of calcite fiber bundles shown in Figures 4b and 5b1, as well as by the different directions of calcite layers in the palisade shown in Figure 7.

When in the liquid state, did these nanospheres contain already formed fibers and liquid material (water + organic substances)? The pore edges in images 8b′, 8c′, and 8d′ indicate that the interior of these hemispheres is filled with fibers whose diameters could not be individualized by SEM. However, considering the hemisphere diameter (57 ± 12) nm and the “clouds” of fibers formed at the edges in Figures 8b′, 8c′, and 8d′, it is inferred that individual fiber diameters may approach the value of the calcite unit cell c-axis (c ∼ 1.7 nm) (47).

The content of the hemispheres shown in Figures 5 and 8 consists of fibers, organic phase, and water. When did fiber nucleation and growth occur—at the moment these nanoparticles adhered to pre-existing lamellae? Once adhered to the surface, surface energy may be sufficient for heterogeneous nucleation (48) and fiber growth. However, the simultaneous coalescence of many droplets should result in fiber misalignment, as suggested by the scheme shown at the top of Figure 8.

The biomineralization rate of laying hens seems to occur more rapidly during calcite crystallization and palisade (PL) growth. This stage until laying is about 8–10 hours (49) to form a layer corresponding to about 80% of eggshell thickness. Another ∼8 hours were spent forming the mammillary layer, whose thickness represents around 10% of eggshell thickness.

#### 3.1.3. Mammillae and Inner Membrane

The images in Figure 9 show the mammillary region using SE and BSE sensors. They correspond to exactly the same areas. The difference is that the BSE sensor captured the presence of a film enveloping the basal plate of the mammillae (9a1). In 9b (SE, topography), we observe the growth of superposed CaCO₃ lamellae (see the semi-quantitative composition of the area by EDS in 9c) starting on the inner face of the membrane that surrounds the egg white. The same image acquired with the BSE sensor (9b1) shows that particles grow within the organic phase. A blue rectangle marks the same area in both images b and b1. Within the rectangle, the lamellae present on the surface of image 9b arise from particles that nucleated on the membrane surface. Sulfur (S) and calcium (Ca) EDS maps are shown in images c1 and c2. Sulfated organic compounds (50,51) coat the collagen membrane.

**Figure 9.**
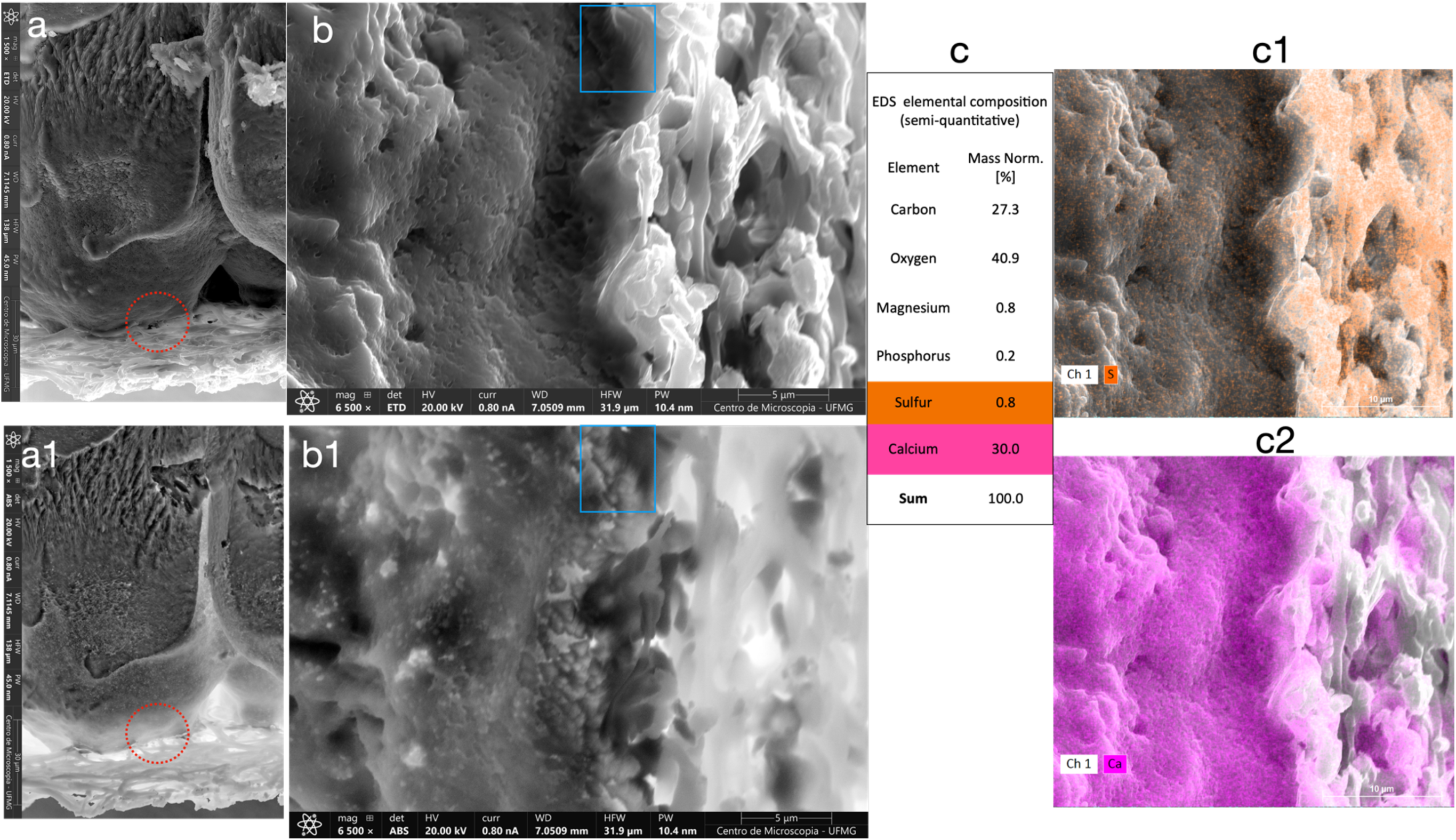
Sequence of SEM images of the same area at the base (13) of the eggshell using SE (9a and 9b) and BSE (9a′ and 9b′) detectors. Comparing image 9a (membrane, basal plate, and mammillae) with 9a1 (BSE), it is possible to observe the existence of a layer surrounding the mammillae. This film shown in a1 is composed of lighter chemical elements (lighter elements appearing in light color), possibly an organic layer, which seems to shape the mammillae and separate them, creating space for the vertical channels/pores that cross the shell. A red dotted circle indicates the area presented in 9b and 9b′. In the topographic image (b), layers growing from the membrane (left side of image b) can be observed, whereas in image b1 particles are visible within the film. A blue rectangle shows that the lamellae present on the surface originate from particles that nucleated on the membrane. The table of elements present in this area (EDS) shows the presence of sulfur-containing compounds (S) (9c1) coating the membrane (50) and the nucleation of CaCO₃ (9c2) growing over the entire outer area of the inner eggshell membrane.

Regarding the site where calcite biomineralization begins, image 9b1 and the calcium map (EDS) in 9c2 confirm what several previous and current studies have indicated: calcite crystallization occurs even within the inner membrane (4,13,52). There, in the inner membrane, the unidimensional (fibrous) morphology of calcite crystals can also be visualized. We did not locate, in the mammillary region, the cubic calcite morphology reported by (53).

#### 3.1.4. Relationship Between Diameters of Cuticle Nanospheres and VL/PL Hemispheres

Figure 10 shows the box plot of the data in Table 3. Different lowercase letters above the boxes indicate statistically significant differences among layers according to the Kruskal–Wallis test (H = 127.08; p = 1.64 × 10⁻²⁶) followed by Dunn’s post-test with Bonferroni correction (p < 0.05). Cuticle nanospheres and hemispheres from the vertical layer (VL) do not differ statistically. Mean diameters of cuticle spheres and VL and PL hemispheres decrease from the cuticle toward the lamella thickness. Spheres forming the pore plug (Figure 3) are larger than the other forms, possibly due to the presence of organic material increasing their diameter.

**Figure 10.**
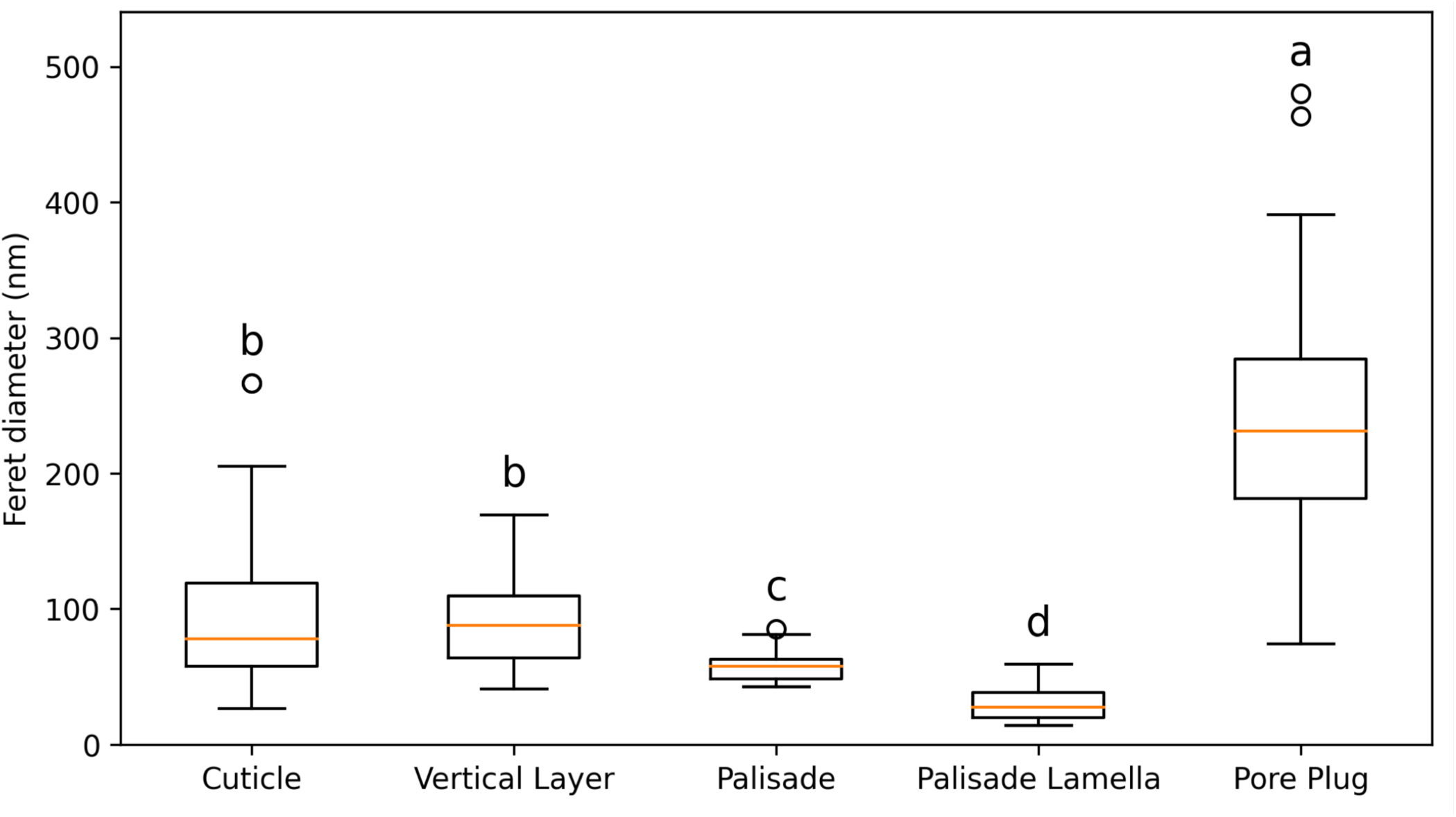
Box plot of the Feret diameter of spherical and hemispherical structures obtained from SEM images (SE mode, topography) in the cuticle (spherical), vertical layer (VL) (hemispherical), and palisade (PL) (hemispherical). *Pore plug* refers to the diameter of spheres present within the pore plug, and *palisade lamella* refers to the thickness of the individual lamellae that constitute the palisade layer. Different lowercase letters above the boxes denote statistically significant differences among layers according to the Kruskal–Wallis test followed by Dunn’s post hoc test with Bonferroni correction (p < 0.05).

### 3.2 Raman Spectroscopy

The Raman spectra (Witec spectrometer) focused on the cuticle region are presented in Figures 11, 12, and 13. Beginning with Figure 11a, which shows an image of a cross section of the shell embedded in resin, where the cuticle region is circled in red. Subsequently, Figure 11b presents the analyzed points. The spectrograms of the four points marked in 11b (1–red, 2–dark blue, 3–green, and 4–light blue) are shown in 11c.

**Figure 11.**
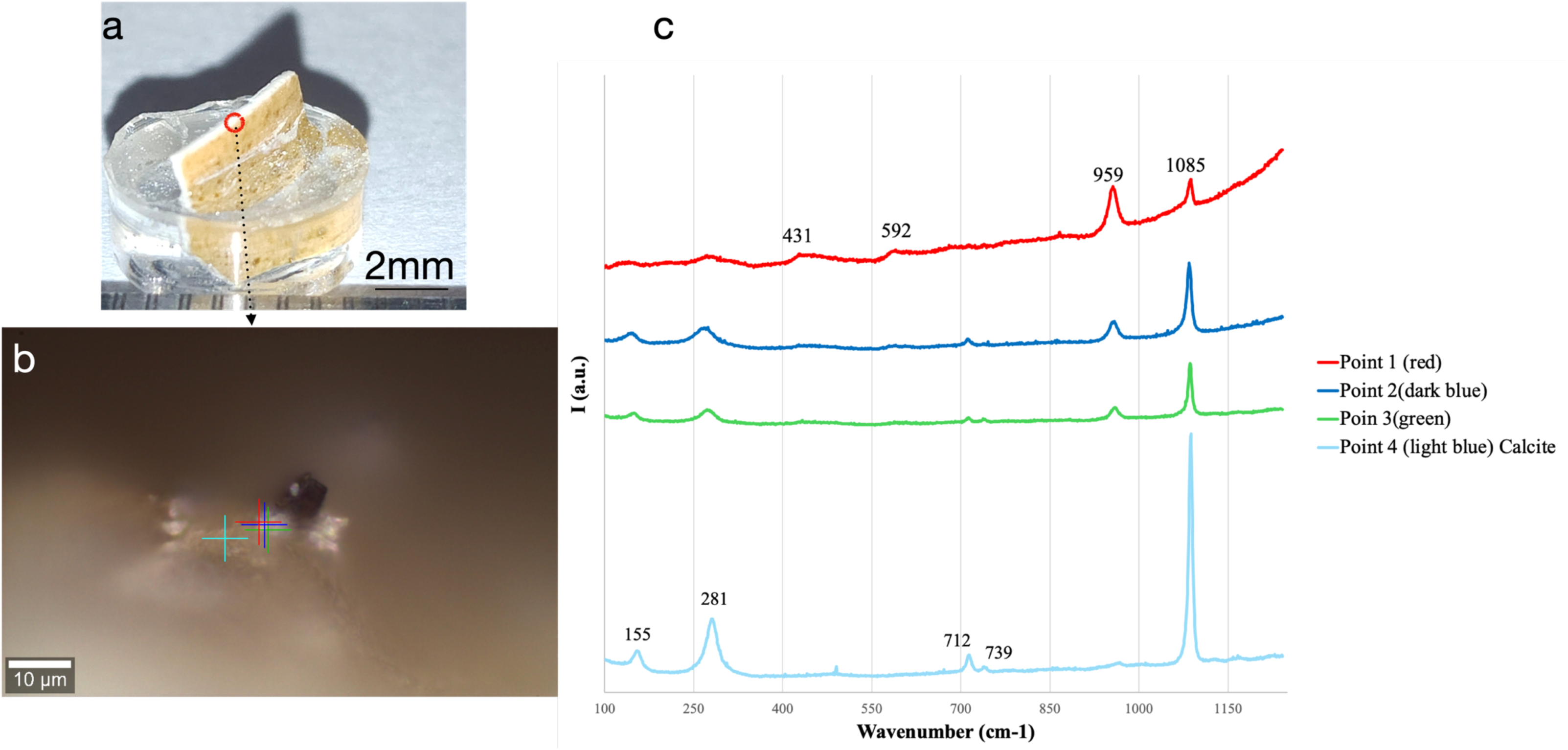
(a) Eggshell sample embedded in resin, with the thickness highlighted by a red circle, for investigation of the cuticle region (brown side)–vertical layer (VL) interface 530 by micro-Raman spectroscopy; (b) Four points marked in the image—red (1), dark blue (2), green (3), and light blue (4)—correspond to the locations investigated in the cuticle (C)–vertical layer (VL) region, and their Raman spectra are presented in (c).

In the spectrum from point 3 (green) to point 1 (red), there is an increase in the intensity of the 959 cm⁻¹ peak, the most intense peak of carbonated hydroxyapatite (CHAp) (54). The other CHAp peaks are also present, especially the 1085 cm⁻¹ peak, indicative that the CHAp is B-type, that is, the apatite is carbonated by substitution of the PO₄³⁻ group by CO₃²⁻ (54,55). CHAp-B is present in vertebrate bones. To the left of these three points is point 4 (light blue spectrum). The main peaks marked (155, 281, 712, 739, and 1085 cm⁻¹, the most intense) correspond to calcite. From the position of the points and spectra, a gradual transition from calcite (Vertical Layer, VL) to CHAp-B is observed. The spectra of points 3 (green) and 2 (dark blue) even indicate the presence of peaks at 155 and 281 cm⁻¹, which decrease in intensity as the main CHAp-B peaks (959 and 1085 cm⁻¹) increase. In the spectrum of point 1 (red), these peaks disappear, and only CHAp-B peaks are observed.

As already documented in the SEM images—especially in Figures 2.b3 and 4b (showing the cuticle–vertical layer VL region)—and indicated by the micro-Raman spectra in Figure 11c, the CHAp-B layer—does not abruptly terminate the calcite VL layer. Rather, an interaction between the cuticle and the VL is observed, indicating a continuous transition from CHAp-B to calcite.

In another experiment (Figure 12), performed on a different Raman instrument (Horiba) to identify the inorganic compounds in the cuticle, two distinct preparations were produced. The spectrum in Figure 12a, obtained at the red The spectrum on the right was acquired at the point marked with a red cross; the spectrum (0–3000 cm⁻¹) corresponds to B-type carbonated hydroxyapatite (54).

**Figure 12.**
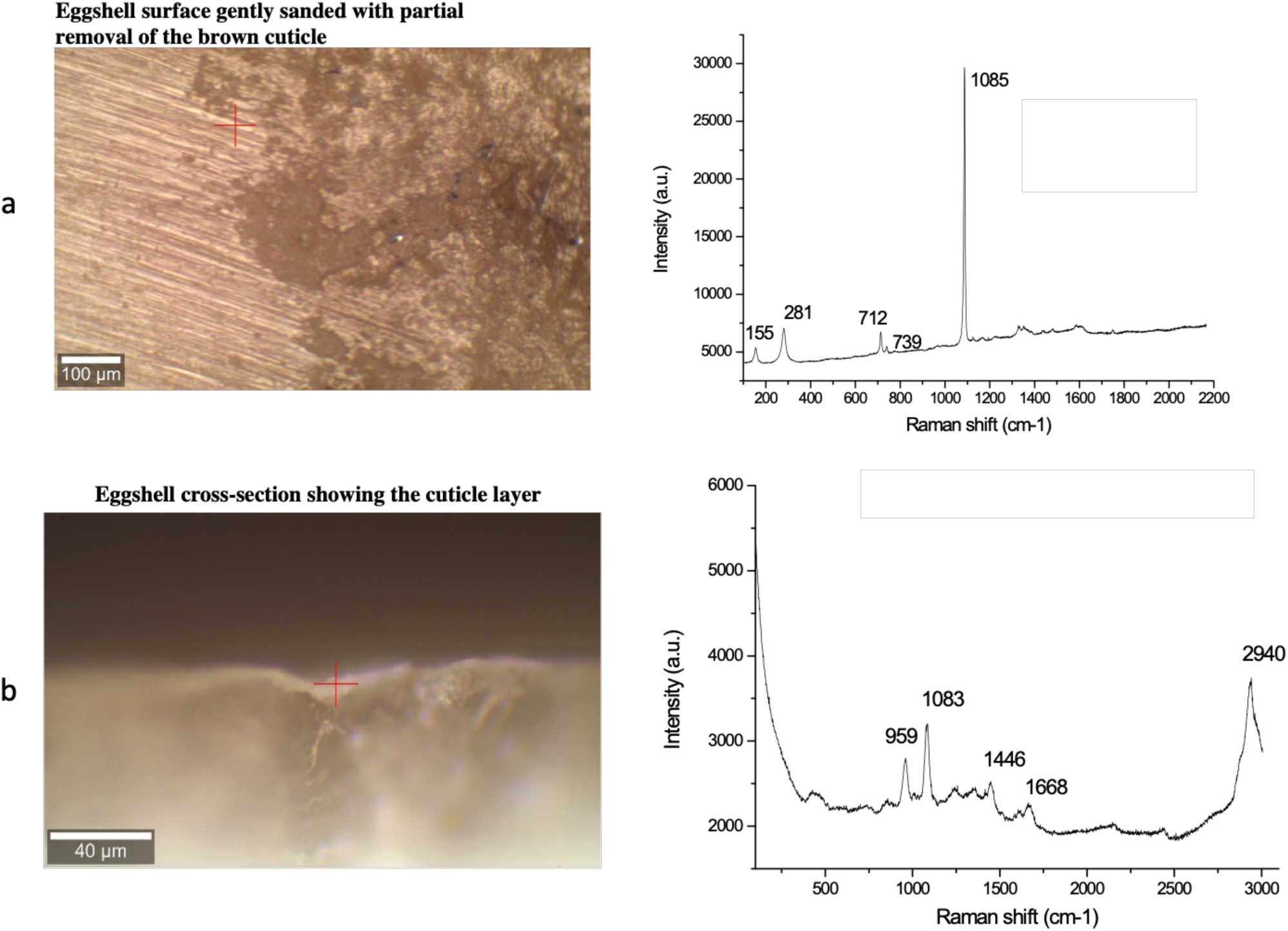
Micro-Raman analysis of two distinct sample preparations: (a) eggshell surface gently sanded with partial removal of the brown cuticle. The spectrum on the right was acquired at the point marked with a red cross; the spectrum corresponds to calcite. (b) Cross-sectional image of the eggshell in the cuticle region.

cross, shows the shell surface with the cuticle (brown) sanded until the calcite layer (white) appeared. Figure 12b shows the Raman spectrum of the cross section of the eggshell (red cross in image 12b) sample. The spectrum in 12b corresponds to carbonated hydroxyapatite CHAp-B.

Figure 13 presents Raman spectra (Horiba) of the external (black) and internal (red) surfaces of a chicken femur fragment (see Materials and Methods). The spectra are identical to each other and indicate CHAp-B as the biomineralized phase. The spectrum is identical to that observed in the eggshell cuticle (light blue line).

**Figure 13.**
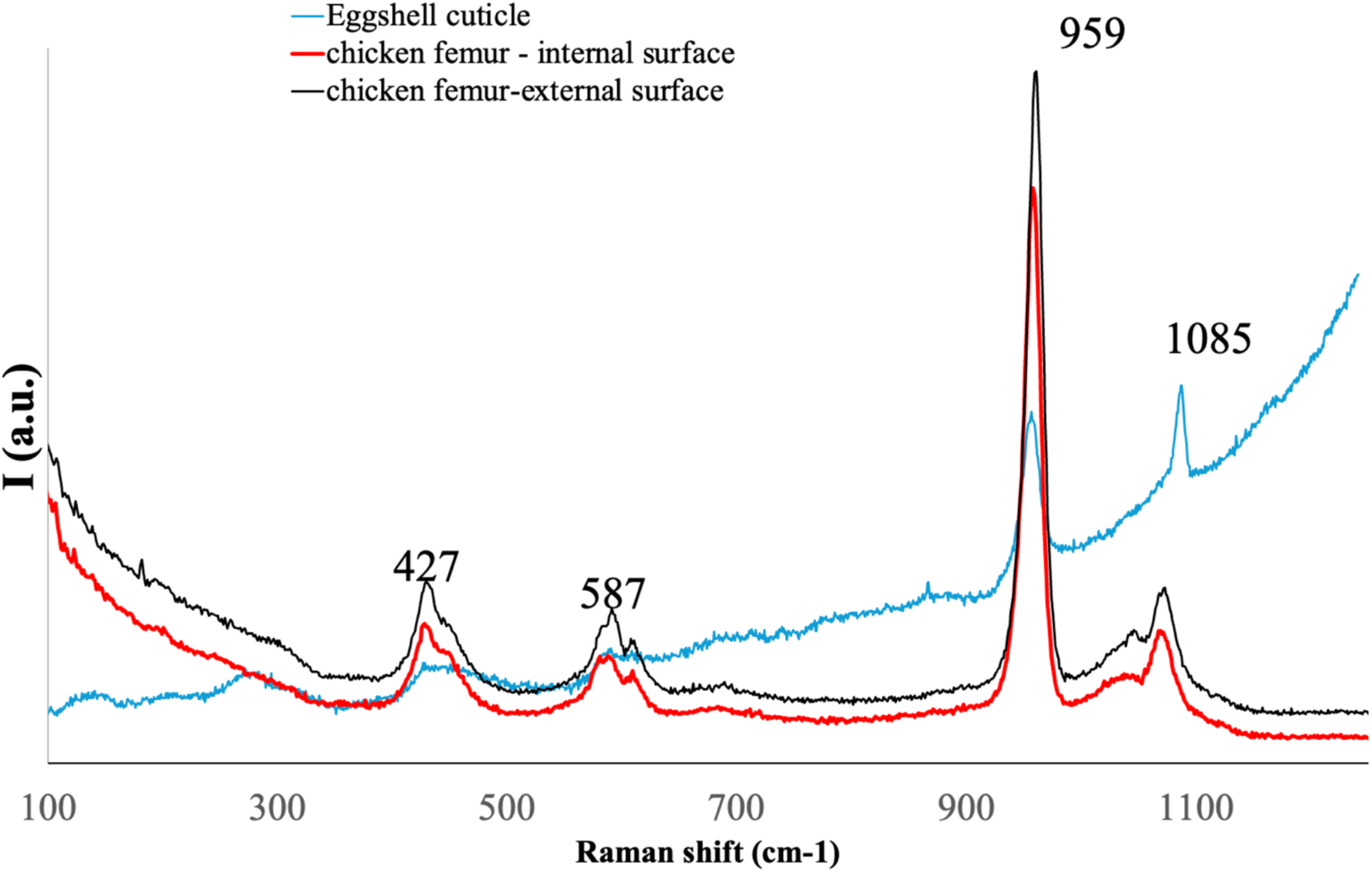
Raman spectra of the external (black line) and internal (red line) surfaces of chicken femur bone and of the eggshell cuticle (light blue 559 line). All three exhibit the main characteristic peaks of B-type hydroxyapatite.

### 3.3 Infrared Spectroscopy (FTIR-ATR)

The spectra obtained using FTIR-ATR coupled to an optical microscope (micro-FTIR) in Figure 14a, recorded in the 700–1700 cm⁻¹ range, compare the cuticle profile (red line) with the spectrum (black line) obtained from the internal surface of a chicken femur fragment (see item 2.1.3 in Materials and Methods). The absorption band occurring in both samples (1026 cm⁻¹ for femur and 1033 cm⁻¹ for cuticle, marked in the spectrum) corresponds to the main hydroxyapatite band. Some studies suggest the band in the interval 1000–1100 cm⁻¹ (54,56,57).

**Figure 14.**
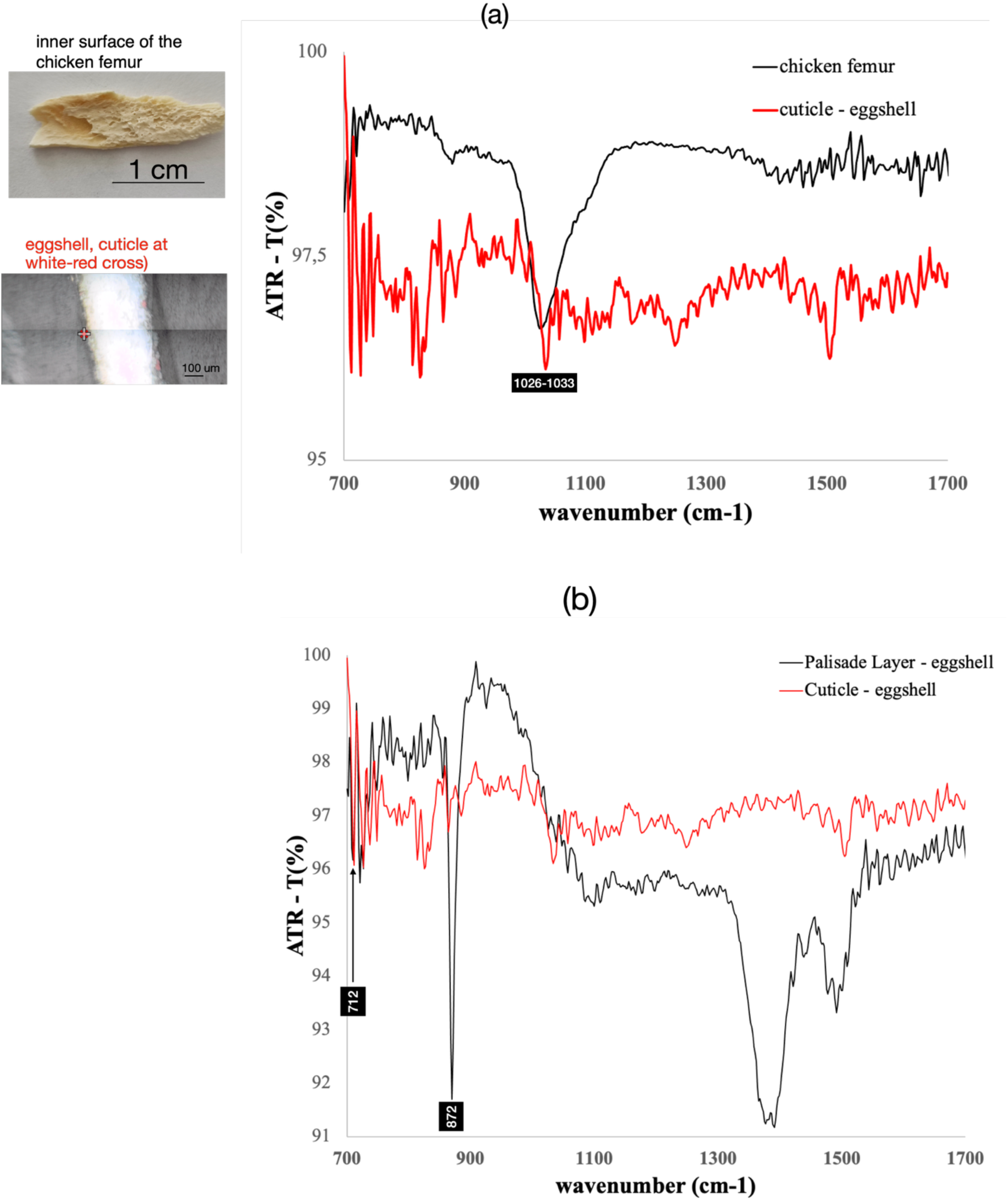
ATR-FTIR spectra (700–1700 cm⁻¹, % transmittance) of: (a) the inner surface of the chicken femur and the eggshell cuticle (see images on the left); (b) the eggshell palisade layer and the cuticle. In (a), the main hydroxyapatite ab-sorption band (1026–1033 cm⁻¹) is observed in both the femur bone and the cuticle. In (b), the two characteristic calcite bands (712 and 872 cm⁻¹) are clearly present in the palisade layer and are much less intense in the cuticle.

Figure 14b shows the spectrum obtained in the palisade layer (PL) region (black line) and that of the cuticle (red line). The PL spectrum corresponds to calcite, with the two main peaks (712 and 872 cm⁻¹, marked in Figure 14b) intense and well defined (58,59). The cuticle spectrum also presents these peaks (712 and 872 cm⁻¹), but with lower intensity, since the main absorption band is at 1033 cm⁻¹, attributed to hydroxyapatite.

Figures S4 and S5 (Supplementary Material) show FTIR spectra of eggshell samples previously thermally treated at 950 °C, exhibiting an absorption band between 1000–1100 cm⁻¹ centered at ∼1033 cm⁻¹, which coincides with the main hydroxyapatite band (56,57). Calcination of calcite at 950 °C greatly reduces the intensity of the main calcium carbonate peaks (712, 872, and 1407 cm⁻¹), facilitating visualization of the main hydroxyapatite absorption band. Anhydrous calcium oxide (CaO) (60), resulting from calcite calcination, presents bands only below 400 cm⁻¹.

## 4. Discussion and Conclusions

A detailed analysis of the layers of the laying hen eggshell was carried out by SEM. Special attention was given to comparing topographic images (SE sensor) with images obtained using the BSE sensor, which reaches depths of tens of micrometers and differentiates, by tonal variations, the chemical composition of the scene. The semi-quantitative values of this composition are provided by EDS spectroscopy (61) coupled to SEM. Other spectroscopic techniques with micrometer resolution (micro-Raman and micro-FTIR) were essential to investigate the region where the cuticle and the vertical layer (VL) come into contact.

In this way, we were able to observe that:

1- Well-defined spheres are present in the cuticle containing phosphorus (P) compounds, with larger diameters in the upper part and smaller diameters in the lower part. Especially in these spheres of the lower layer, which interact with the VL, the P signal is higher. Viewed in detail, these spherical nanovesicles contain, within their interior, aggregates where the P compound is located;

2- Cuticle–Vertical Layer (VL): No boundary was observed between the base of the cuticle and the calcite vertical layer (VL). A clear interaction was observed with the contents of the cuticle vesicles. The interaction is mechanically demonstrated by fracture that occurs in the VL but keeps the VL adhered to the cuticle intact (Figure 2). In BSE images it became evident that the VL is composed of fibers with diameters of a few nanometers (Figure 4). The fibrous morphology of calcite is known in calcite biomineralization processes (62–64);

3- Vertical Layer: SEM SE (topography) images visualize the VL as lamellae that grow by adhesion of nanospheres (Figures 5a and 5b). In the BSE image, the lamellae are composed of fibers that grow through interaction with the cuticle. Bubble-shaped pores are present in the VL and show the same morphology as the pores of the palisade layer (PL);

4- Palisade (PL): the palisade, which accounts for ∼80% of the shell thickness, is also composed of nanometric-thickness lamellae. In the palisade, the calcite lamella is the smallest unit discernible by SEM in both SE (topography) and BSE images (Figure 6). The lamellae are not morphologically homogeneous. There are lighter regions, with higher concentration of calcite fibers. As already reported in the literature, we observed high concentration of “bubble-type” pores (and of similar diameter) throughout the palisade. These “bubble-type” pores may represent more than 20% of the shell volume.

Notable in the palisade is the presence of nano-hemispheres adhered to the lamellae and in the pore region. Images 5a and 5b show the lamella growth process by adhesion and coalescence of nanospheres (or nanodroplets). In images 5d1, 8b′, 8c′ and 8d′ (BSE images), it is possible to discern, at the pore contour, the presence of calcite fibers.

In the upper right corner of Figure 8, the growth process of the palisade and the vertical layer by adhesion of nanodroplets containing already formed fibers or fibers that form, by heterogeneous nucleation, upon adhesion to the existing surface is suggested. The fibers should not have very long length, if already formed within the nanodroplet, since misalignment would occur between fibers of nanodroplets that coalesce.

5- By size (Feret diameter), there is statistical correlation between the cuticle nanospheres and the nano-hemispheres of the vertical layer VL (Figure 10 and Table 3). The hemispheres of the palisade are even smaller, and even smaller is the thickness of the calcite lamellae measured in the palisade. These differences must be attributed to greater hydration of the nanodroplets in the cuticle compared to the hemispheres of the palisade and to the contraction that results in the formation of calcite lamellae. Therefore, it is considered that the phosphate nanospheres correlate with the hemispheres observed in the vertical layer and in the palisade. The hemispheres observed in the VL and PL are the units responsible for shell growth by additive deposition.

6- Prior to solidification, these nanodroplets are presumed to have carried water, the mineral phase, and the organic components constituting the ultrastructure. Everything would be formed at once. The pres-ence of nanodroplets in the completed shell provides evidence of the high rate of eggshell biominerali-zation, since nano-hemispheres represent an incomplete biomineralization stage that did not fully trans-form into lamellae.

If this is a generalized phenomenon in laying hens subjected to high productivity demands, this may influence the mechanical resistance of the eggshell.

7- Calcite biomineralization in the mammilla/membrane region of the inner eggshell (Figures 9b and 9b1) appears to occur within a viscous phase, which is the uterine environment. Calcite crystals appear embedded in the organic matrix. In 9.c2 it is possible to observe that calcite occurs even within the inner eggshell membrane.

8- Raman spectra in the cuticle indicated the presence of B-type carbonated hydroxyapatite (CHAp-B), the same phase present in bones. Spectra obtained at micrometric points placed along a line perpendicular to the cuticle–vertical layer (VL) interface indicate continuity, with CHAp-B peaks decreasing in intensity as the points approach the vertical layer (Figure 11), until CHAp-B reduces to a minimum to give place to calcite (Figure 11.c).

To confirm the presence of this type of hydroxyapatite, another experiment, using another micro-Raman spectrometer (Figure 12), indicated the CHAp-B profile in a micrometric area of the cuticle.

9- Raman spectra of CHAp-B, present on the internal (including medullary bone) and external surfaces of chicken femur, coincide with the spectrum of CHAp-B present in the eggshell cuticle (Figure 13).

10- The vesicle model carrying the mineral phase in its interior appears consistent (43). However, given the presence of CHAp-B in the cuticle, it is plausible—though not supported by the experimental data of this study—that hydroxyapatite (amorphous or crystalline) is conveyed to the shell through vesicular transport. To detect phosphorus by SEM-EDS, it is necessary to review the experimental procedure, as demonstrated by the tests presented in Figure S6 of the Supplementary Material. Another plausible ex-planation for vesicular transport of hydroxyapatite is metabolic economy. CHAp-B, mobilized from the hen’s medullary bone, could reach the uterus via the bloodstream without the need for additional trans-formation or energetic cost.

11- Although the experimental data of this study are not exhaustive or definitive, the results suggest that the conversion of CHAp-B into CaCO₃ may facilitate the release of metabolically available phosphate for the bird.

12- Embryonic skeleton growth is, at least in the first stage of its development (65,66), associated with the phosphate present in the egg yolk. The complex structure of the chorioallantoic membrane (CAM) (67–69) and its interaction with the eggshell is not the subject of this work. In the second stage of chick skeleton growth (65), absorption occurs, through the CAM circulatory system, of the calcium reserve body (CRB) located at the base of the mammillae and of the calcite in the mammillae. The transformation of calcite into hydroxyapatite is energetically simpler, as shown by dozens of studies that use eggshell as raw material for HAP production (70) and studies using the technique of in ovo injection (IOI)(71). Calcite, in the form of nanofibers, should greatly facilitate this calcite → CHAp-B reaction.

## 5. ACKNOWLEDGEMENTS

The author AVC would like to thank FAPEMIG- Fundação de Amparo à Pesquisa do Estado de Minas Gerais (Grant APQ-04741-24 and BIP-00140-23) and National Council for Scientific and Technological Development (CNPq), Brazil (Grant 404383/2023-8) for the financial support. The authors are grateful to Professor Isolda Maria de Castro Mendes (Chemistry/UFMG) for preparing the samples used in the Raman spectroscopy analyses presented in Figures 11 and 12. APG acknowledges the support of Sis-NANO, FAPEMIG, FINEP, and LCPNano.

## 6. SUPPLEMENTARY MATERIALS

Supplementary material associated with this article is available in the online version of this preprint.

## 7. Author Contributions

Conceptualization, methodology and writing, review and editing, AVC ; SEM-EDS experiments, RNF; Raman experiments and data analysis validation, MSD and APG; FTIR-ATR, XRD, TGA experiments and data analysis validation, LHRS and LLA. All authors have read and agreed to the published version of the manuscript.

## 8. Data Availability Statement

The data supporting the results of this article will be made available by the authors on request.

