## Supplemental Figs S1 to S7 for "Phosphate and Carbonate in the Biomineralization of Chicken Eggshells and the Increase in Eggshell Thickness through Nanodroplet Addition"

### 1- X-ray diffraction (XRD)

X-ray diffraction (XRD) of eggshell samples thermally treated in air at temperatures of 150, 350, 550, 750, and 950 °C. Up to 750 °C, the main thermodynamic phase is calcite ($CaCO_3$; main peaks at 29.4°, 23.0°, and 39.4° 2θ). Above this temperature, a $CO_2$ volatilization process occurs, and at 950 °C the main phase present is calcium oxide (CaO; main peaks at 37.2°, 32.2°, and 53.8° 2θ), as shown in Figure S1. However, $CaCO_3$ peaks are still detectable, although with markedly reduced intensity.

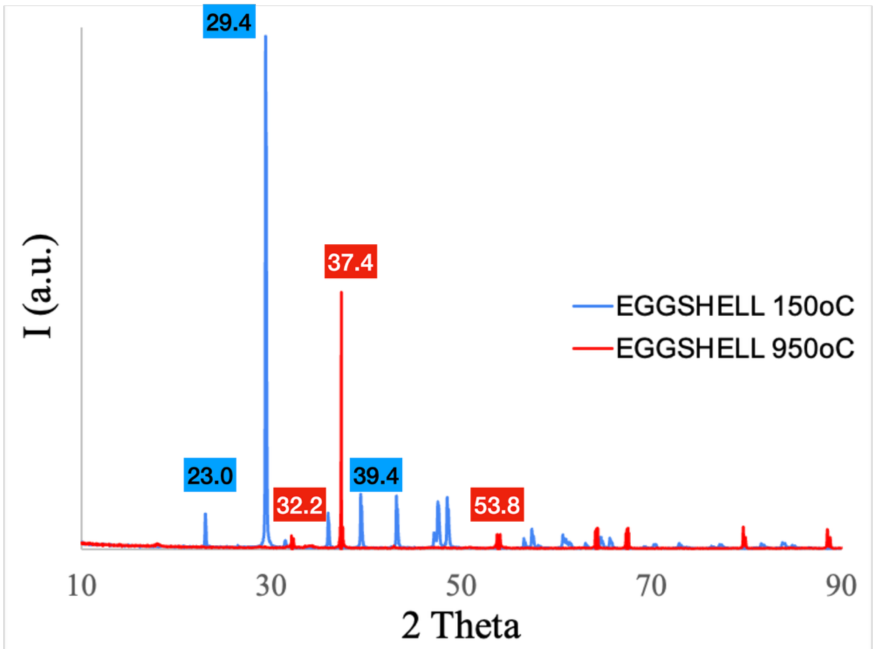


**Figure S1 –** X-ray diffraction (XRD) of eggshells thermally treated in air at 150 °C (blue) and 950 °C (red) for 1 hour. The phase at 150 °C is calcite ($CaCO_3$, blue peaks), whereas at 950 °C the phases are calcium oxide (CaO, red peaks) and calcite.

### 2- Thermogravimetric Analysis (TGA)

Thermogravimetric analysis (TGA) of eggshell samples previously thermally treated in air at 150, 350, 550, 750, and 950 °C. The tested samples had different masses. To facilitate analysis in a single graph (Figure S2), the experimental curves were adjusted for comparison.

The sample initially treated at 150 °C (orange line, Figure S2) still shows some mass loss related to water, followed by additional loss of other volatiles from 250 °C to 600 °C. The sample treated at 350 °C also exhibits mass loss, but starting at 380 °C. The CaO sample, obtained by calcination of the eggshell at 950 °C (green line), still presents mass loss in the 350–450 °C range, possibly due to adsorption of atmospheric gases.

A $CaCO_3$ sample was also analyzed for comparison. Unlike the eggshell samples, $CaCO_3$ does not exhibit mass loss within the analyzed temperature range (up to 600 °C).

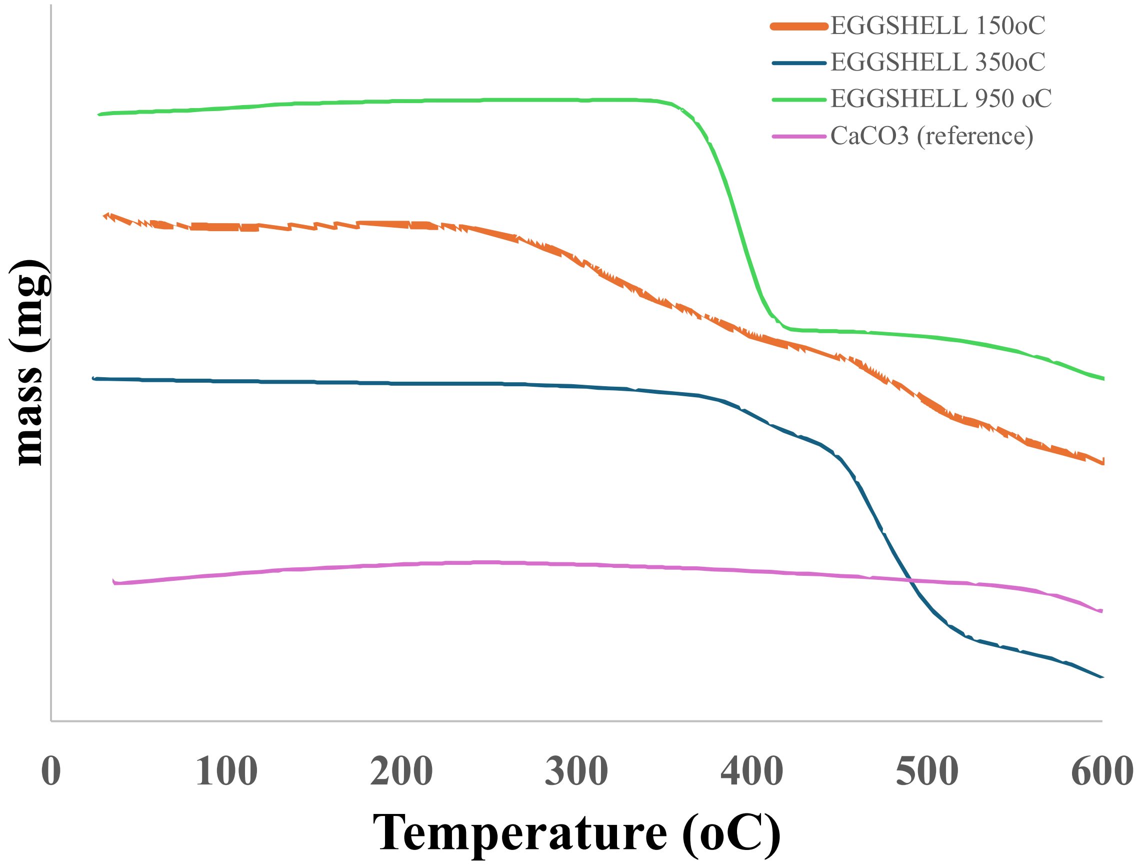


**Figure S2** – Thermogravimetric analysis of eggshell samples previously treated at 150, 350, and 950 °C. For comparison, $CaCO_3$ was also analyzed. The tests were performed up to 600 °C with a heating rate of 10 °C/min.

#### 3 – Infrared Spectroscopy (FTIR)

Figure S3 presents the infrared (FTIR) spectra of eggshell samples thermally treated at 150, 350, 550, 750, and 950 °C. The $CaCO_3$ absorption bands centered at 1407, 872, and 712 $cm^{-1}$ progressively decrease in intensity as the thermal treatment temperature increases. In the sample treated at 950 °C, the bands still present coincide with the $CaCO_3$ absorption bands (3–5). The CaO absorption bands would be located between 250–400 $cm^{-1}$ .

It is important to note that an absorption band located between 1030–1090 $cm^{-1}$ (sample treated at 150 °C) becomes broader and more intense in the sample treated at 950 °C.

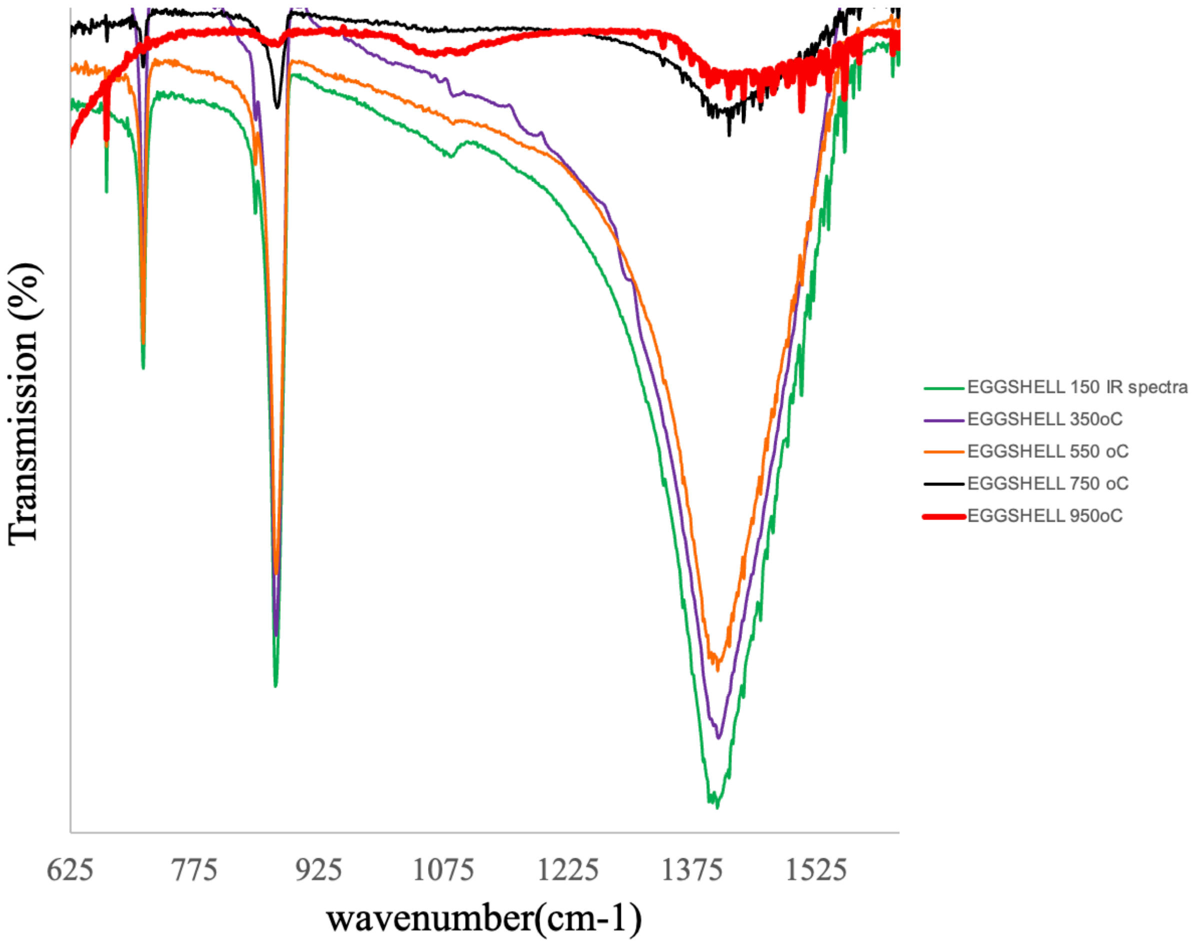


**Figure S3** – Infrared (FTIR) spectrum of eggshell samples previously treated at 150, 350, 550, 750, and 950 °C. The bands of samples treated up to 750 °C correspond to $CaCO_3$ and remain present, although much less intense, in the sample treated at 950 °C. In this sample (red line), the absorption band in the 1000–1100 $cm^{-1}$ range (centered at ~1033 $cm^{-1}$) is not attributed to $CaCO_3$ (6,7).

Figure S4 compares the spectrum of the eggshell sample treated at 950 °C with the spectrum of a bovine hydroxyapatite sample (see item 2.1.4 in Materials and Methods). The 1000–1100 cm$^{-1}$ absorption band of this sample (red curve) partially coincides with the absorption band attributed to the phosphate group of bovine hydroxyapatite (blue curve), between 900–1100 cm$^{-1}$, with peaks at 960 and 1089 cm$^{-1}$ and centered at ~1026 cm$^{-1}$. This band is not attributed to calcium oxide (CaO), which, in its anhydrous form, exhibits absorption bands below 400 cm$^{-1}$ (8).

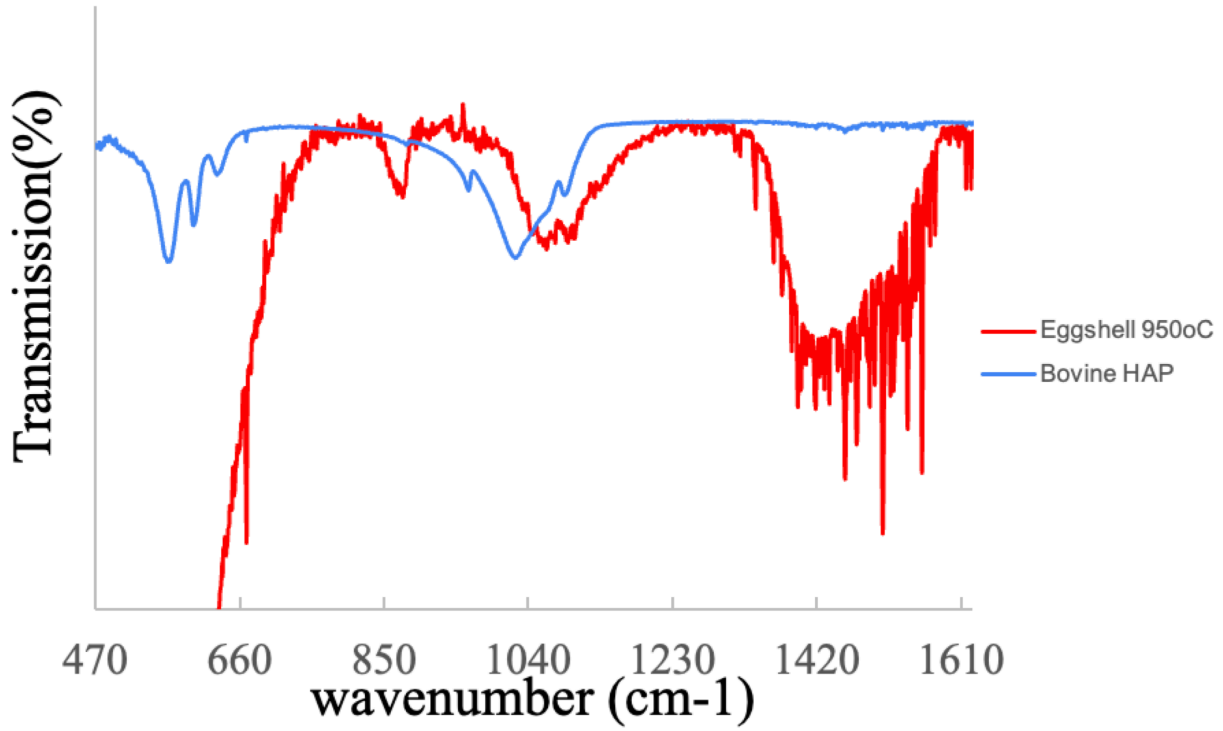


**Figure S4** – Infrared (FTIR) spectra: (i) of an eggshell sample previously treated at 950 °C (red curve) and (ii) of a bovine hydroxyapatite sample. The spectrum of the sample treated at 950 °C still exhibits bands of residual $CaCO_3$; however, the band between 1000–1100 cm$^{-1}$ partially coincides with the 900–1100 cm$^{-1}$ band of bovine hydroxyapatite.

### 4 – Identification of phosphorus (P) by EDS using carbon (C) or gold (Au) coating

In the preparation of non-conductive samples for SEM, it is well recognized that gold (Au) coating interferes with phosphorus (P) identification and its quantification by EDS.

Figure S5 presents the EDS spectroscopy results in the cuticle region of eggshells from two distinct areas of a gold-coated sample (solid red and blue lines) and from three distinct areas of the cuticle in a carbon-coated sample (dashed green, black, and purple lines). As observed in the spectra in Figure S5, the X-ray emission peak of gold (Mα, 2.12 keV) overlaps with the phosphorus peak

(Kα, 2.03 keV). In the carbon-coated sample, the phosphorus emission peak is not affected. Therefore, SEM-EDS studies using gold coating on eggshell samples may underestimate the concentration of phosphorus and phosphate present in the sample.

Dennis et al. (9), in 1996, reported the presence of hydroxyapatite in the cuticle because they used palladium (Pd) coating on the sample. The Pd emission peaks are 2.84 keV (Lα) and 2.99 keV (Lβ).

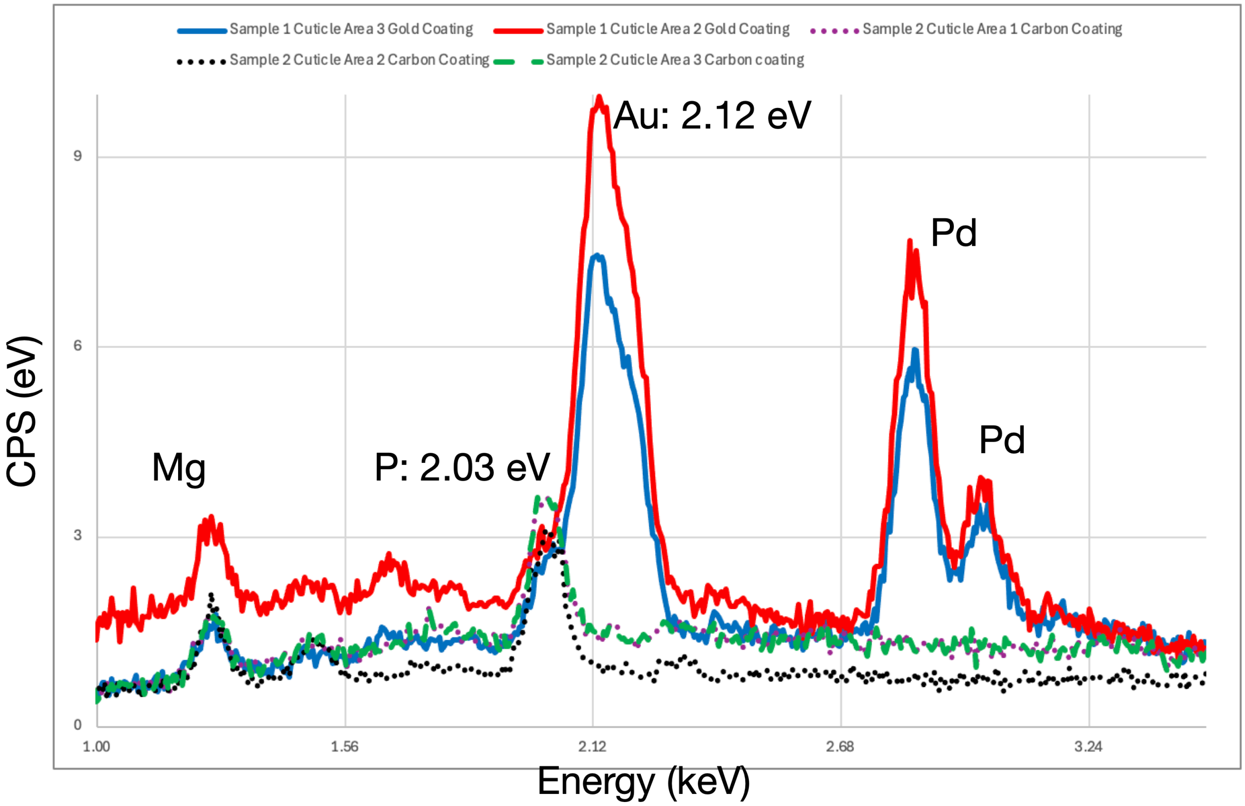


**Figure S5 –** X-ray emission peaks obtained by EDS spectroscopy in the cuticle region from two distinct areas of a gold–palladium-coated eggshell sample (solid red and blue lines). Three dashed lines (black, purple, and green) correspond to a carbon-coated eggshell sample. The gold emission peak (2.12 keV) overlaps with the phosphorus peak (2.03 keV).

**5 – Structure of the Mammillary Base**

As the shell thickness increases, the successive calcite layers that nucleated and grew from the inner eggshell membrane form structures called mammillae and, a few micrometers above, the palisade layer develops. The SEM image shown in Figure S6 displays a structure forming a kind of calcite crown at the tip of a mammilla (9). This crown is marked with number 3 in the image.

Number 1 indicates the region called the Calcium Reserve Body (CRB). In this region, spheres with diameters < 1 micrometer can be observed. These spheres would contain the mineral phase that is absorbed by the blood circulation present in the chorioallantoic membrane (CAM) (10). The length (white dotted line) was evaluated as (17.2 ± 2.4) µm.

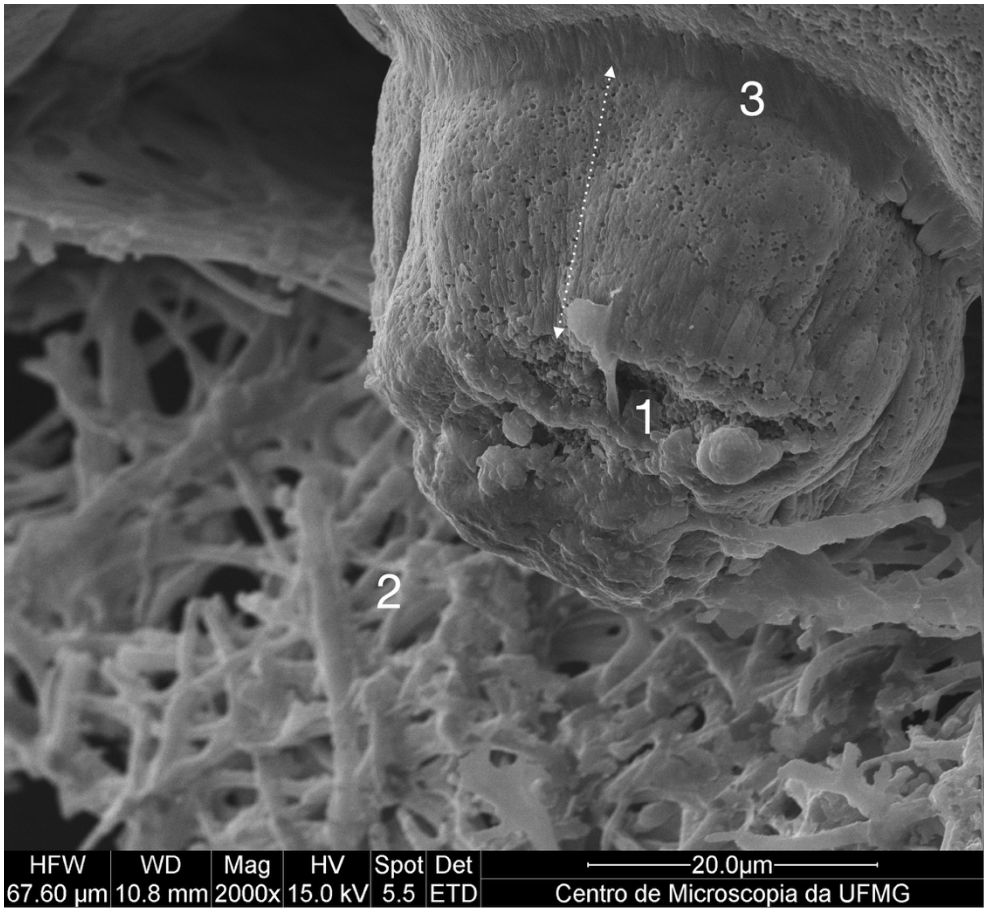


**Figure S6 –** Mammillary region that, upon detachment from the inner membrane (marked with number 2) of the eggshell, reveals the interior of the Calcium Reserve Body (CRB), marked with number 1. Spheres with diameters < 1 µm constitute this reserve. The crown, marked with number 3, indicates the beginning of the palisade layer (PL). The length (dotted line) was evaluated as (17.2 ± 2.4) µm.

**6 – X-ray Diffraction (XRD) of Bovine Hydroxyapatite**

As described in item 2.1.4 of the Materials and Methods section of this work, after obtaining the hydroxyapatite sample derived from bovine bones, X-ray diffraction was performed. The result is presented in Figure S7, where both the reference material and the sample indicate that they are hydroxyapatite.

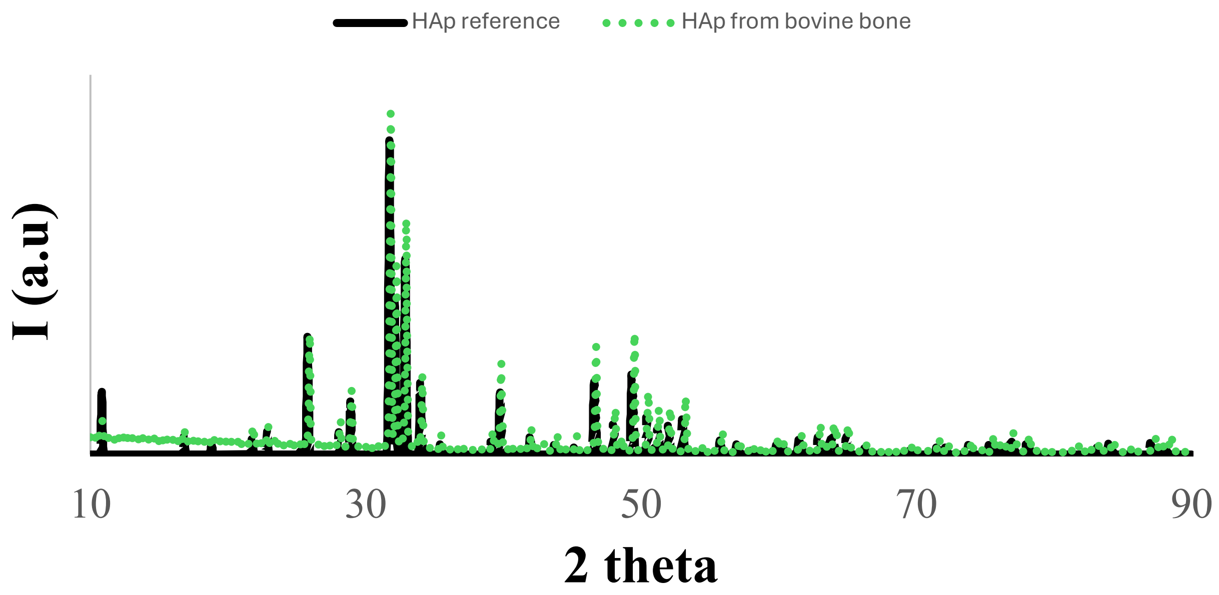


**Figure S7 –** XRD spectrum of the reference hydroxyapatite material (black line) and the sample obtained from bovine bone (green dotted line) as described in item 2.1.4 of the Materials and Methods section of this work.
